# RAS activation via CRLF2 signaling is a widespread mechanism in Down syndrome acute lymphoblastic leukemia regardless of RAS mutations

**DOI:** 10.1101/2020.02.03.931725

**Authors:** David Koschut, Debleena Ray, Zhenhua Li, Emanuela Giarin, Jürgen Groet, Ivan Alić, Shirley Kow-Yin Kham, Wee Joo Chng, Hany Ariffin, David M. Weinstock, Allen Eng-Juh Yeoh, Giuseppe Basso, Dean Nižetić

## Abstract

**Background:** Down syndrome acute lymphoblastic leukemia (DS-ALL) is characterized by the high frequency of CRLF2-rearrangements, JAK2-mutations, or RAS-pathway mutations. Intriguingly, JAK2 and RAS mutations are mutually exclusive in leukemic sub-clones, causing dichotomy in therapeutic target choices.

**Results:** Here we show that in primary leukemic cells from DS-ALL, in the absence of RAS-mutations, wild-type (wt)RAS is active, and/or can be induced by the physiological ligand TSLP of the transmembrane-receptor CRLF2. We show active/inducible RAS in 14/20 (70%) of primary DS-ALL samples analyzed, 8 of which had no RAS-mutations, but 75% of those had either mutated or hyperphosphorylated JAK2. No wtRAS cases with mutated/hyperphosphorylated JAK2 were observed that lacked activated RAS protein. We prove in a cell model that elevated CRLF2 in combination with constitutionally active JAK2 is sufficient to activate wtRAS. We show that TSLP boosts the direct binding of active PTPN11 to wtRAS. Pre-inhibition of RAS or PTPN11, but not of PI3K or JAK signaling, prevented TSLP-induced RAS-GTP boost.

Using multivariate-clustering based on RAS-activity/inducibility we show significant separation between standard-risk and high-risk DS-ALL groups. Cox proportional-hazards model showed protein-activity (but not mutation status) as independently predictive of outcome.

**Conclusions:** Our data indicate that RAS protein activity levels (and not JAK2/RAS mutation profiles), are predictive of outcome. Importantly, our data suggest that inhibition of RAS and direct RAS-pathway components should be combined with PI3K/mTOR and/or JAK2 inhibitors for high-risk cases. Therapeutically this is relevant for >75% of DS-ALL and our additional data suggest that it warrants further investigation in high-risk non-DS-ALL.

## Introduction

Acute lymphoblastic leukemia (ALL) is the most common malignancy and cancer-related cause of death at pediatric age(1, 2). Despite a considerable success rate of standard chemotherapy treatments, as many as 10-15% of children with ALL have recurrent disease (relapses)(3, 4). Patients with high-risk (HR) forms of ALL show increased incidence of relapses, poorer prognosis and lower overall 5-year survival rates following relapse(5). Recently, significant progress has been achieved in understanding the mechanistic consequences of individual pathways affected in HR-ALL, and the resulting selection of therapeutic targets leading to clinical trials using pathway-specific drugs, such as JAK/STAT inhibitors(6). Recent detailed studies of the evolution of acquired genomic changes in ALL identified certain sub-types as being particularly HR forms(7, 8). Among these are hypodiploid ALL(9), Philadelphia chromosome-like (Ph-like) type (defined as a type of ALL with the genomic profile similar to that of the Ph+ ALL)(8, 10, 11), ALL with an intrachromosomal amplification of chromosome 21 (iAMP21)(12, 13), and ALL in children with Down syndrome (DS-ALL)(14, 15).

The acquired mutations landscape does not find a unifying profile that distinguishes HR childhood ALL from non-HR childhood ALL, suggesting the need for individualized therapy approach(16) preceded by individual patient sub-type assignment based on the mutational profile analysis. While Ph-like ALL has a high incidence (60%) of genomic rearrangements leading to an increased expression of the receptor to the cytokine TSLP, CRLF2(17), and more than half of these have mutations in JAK and IL7R pathway − including constitutionally activating JAK2 mutations(11, 18, 19), less than 10% of Ph-like ALL also acquire RAS/MAPK pathway mutations. DS-ALL is distinguished by the similarly high presence of both CRLF2-rearrangements (60%) (with JAK2 mutations at 32%), with a higher proportion of RAS-MAPK pathway mutations (36%)(20, 21). Intriguingly, a near complete mutual exclusion between JAK2 and RAS mutations in diagnosis samples, or individual sub-clones of relapse samples of DS-ALL is repeatedly observable(20, 21).

We hypothesized that the reason for this mutual exclusion is that increased CRLF2-levels in combination with JAK2 activation could be sufficient to activate wild-type (wt) RAS protein in the absence of *RAS* mutations.

## Materials and methods

### Antibodies, inhibitors, and cytokines

The primary antibodies against β-actin (Cat.#ab8227; Western blot (WB)1:10,000) and GRB2 (Cat.#ab86713; PLA 1:100) were purchased from Abcam (Cambridge, UK); the antibody against CRLF2 (Cat.#AF981; WB 1 µg/mL) was purchased from R&D Systems (Minneapolis, US). The primary antibodies against pan-RAS (Cat.#8832; WB1:200), phospho-bRAF (Cat.#2696; WB1:1,000, PLA1:100), JAK2 (Cat.#3230; WB1:900), HA-tag (Cat.#2367; WB1:1,000), MEK1/2 (Cat.#8727; WB1:1,000), phospho-MEK1/2 (Cat.#9154; WB1:1,000), RPS6 (Cat.#2317; PLA1:50, WB1:1,000) and phospho-STAT5 (Cat.#4322; WB1:1,000), phospho-JAK2 (Cat.#3771; WB1:1,000), ERK1/2 (Cat.#9102; WB1:1,000), phospho-ERK1/2 (Cat.#9101; WB1:1,500), GRB2 (Cat.#3972; WB1:1,500), phospho-RPS6 (Cat.#2211; PLA1:50, WB1:1,000), PI3Kp110α (Cat.#4249; PLA 1:100), SOS1 (Cat.#5890; PLA 1:100), PTPN11 (Cat.#3752; WB1:1,000, PLA1:100), phospho-PTPN11 (Cat.#3751; WB1:1,000, PLA1:100), and STAT5 (Cat.#9363; WB1:1,000) were purchased from Cell Signaling Technology (Danvers, US). The primary antibodies raised against KRAS (Cat.#sc-30; WB1:180), NRAS (Cat.#sc-31; WB1:160) and HRAS (WB1:170) were bought from Santa Cruz Biotechnology (Dallas, US). The primary antibodies used in immunofluorescence against pan-RAS (Cat.#MA1-012; IF1:100, PLA1:100) and bRAF (Cat.#PA5-14926; IF1:50, PLA1:50, WB1:1,000) were purchased from ThermoFisher Scientific (Waltham, US), as was SOS1 (Cat.#MA5-17234; PLA1:100).

Secondary HRP-conjugated antibodies against mouse (Cat.#ab97023; WB1:8,000), rabbit (Cat.#ab97051; WB1:9,000), or goat (Cat.#ab97100; WB1:7,000) IgG species were obtained from Abcam. The secondary fluorescent antibodies anti-mouse IgG Alexa Fluor 488 (Cat.#A11029) and anti-rabbit IgG Alexa Fluor 594 (Cat.#A11037) were purchased from ThermoFisher Scientific.

The small molecule inhibitors PI-103 (PI3K/mTOR-inh.; Cat.#S1038), Ruxolitinib (JAK-inh.; Cat.#S1378), Salirasib (RAS-inh.; Cat.#S7684), Rigosertib (RAS-signaling-inh.; Cat.# S1362), PD0325901 (MEK1/2-inh.; Cat.#: S7684), and Vemurafenib (RAF-inh.; Cat.#S1267) were purchased from Selleck Chemicals (Houston, US). Additionally, the PTP inhibitor XXXI/II-B08 (PTPN11-inh.; Cat.#565852; EMD Millipore, Burlington, US) was purchased. All inhibitors were reconstituted in dimethyl sulfoxide (DMSO; Cat.#D2650; Sigma-Aldrich, St. Louis, US).

The cytokine used for Ba/F3 culturing was 10 ng/mL murine IL-3 (Cat.#31310-03-10; Gold Biotechnology, St Louis, US).

### Patient samples

Surplus clinical or archived clinical material for peripheral blood/bone marrow samples of DS-ALL and non-DS ALL patients was collected by the tissue bank of the Italian Association for Paediatric Haematology-Oncology. In accordance with the Declaration of Helsinki, informed written consent was obtained by the tissue bank for all subjects. Samples were processed and stored in the tissue bank at The Blizard Institute, which is licensed for tissue storage and monitored by UK-Human Tissue Authority. Detailed clinical description of studied DS-ALLs and Non-DS B-ALLs is available in Supplementary-Tab.S1. Detailed cytogenetics was available in 12 cases.

MS2003/2010 cohort(22) RNA-seq data was submitted to the European Genome-phenome Archive (Accession# EGAS00001001858).

### Cell culture

Ba/F3 (Cat.#RCB0805), a murine IL-3 dependent pro-B cell line, was obtained from RIKEN BioResource Center (Tsukuba, Japan) and MUTZ-5 (Cat.#ACC490), a human B cell precursor leukemia cell line established at relapse, was obtained from the Leibniz Institute DSMZ (Braunschweig, Germany); authenticated via multiplex PCR of minisatellite markers. Mycoplasma-free cells were routinely passaged (passage range for shown experiments: 15-35) according to the respective cell bank recommendations. Handling of primary patient samples is described in detail in Supplementary-Fig.S4A legend.

### RAS activity assays

Cells were left uninduced or induced with human TSLP at 37 °C. Whenever indicated, DMSO or inhibitors were added for 3 hrs before TSLP-induction. Cells were lysed on ice at 1000 cells/µL lysis buffer according to the manufacturer’s protocol of the active RAS detection kit (Cat.#8821; Cell Signaling Technology). Total protein concentrations of samples were measured using a BCA protein-assay kit (Cat.#23225; ThermoFisher Scientific). 50 µg total protein was loaded per column of the active RAS detection kit for WB. In the RAS activation assay kit for ELISA (Cat.#17-497; EMD Millipore), 12 µg total protein was used at 100 ng/µL and the RAS-GTP pull-down was measured using a Synergy H1 plate reader (BioTek, Winooski, US) in luminescent mode.

### SDS-PAGE and WB

Protein lysates (see “RAS activity assays”) were mixed with 4×Laemmli buffer (Cat.#161-0747; Bio-Rad Laboratories, Hercules, US) containing fresh 200 mM DL-Dithiothreitol (Cat.#3483-12-3; Sigma-Aldrich). For the majority of samples 5 µg total protein could be loaded. SDS-PAGE with 11%-resolving/5%-stacking acrylamide gels, and WB on PVDF-membrane (Cat.#88518; ThermoFisher Scientific) were performed according to the standard protocol of the equipment-manufacturer (Bio-Rad Laboratories). Each PVDF-membrane piece (per antibody) was separately imaged via the auto-exposure function of the ChemiDoc-MP imaging system (Bio-Rad Laboratories). Supplementary-Fig.S4B legend describes the WB-signal quantification in detail. PVDF membranes were stripped from antibodies using the Restore WB-Stripping buffer (Cat.#21059; ThermoFisher Scientific).

### Proximity ligation assay (PLA)

MUTZ-5 cells at 1×10^6^ cells/mL density were either not induced or induced with 20 ng/mL TSLP for 10 min. Where indicated, cells were pre-treated with either DMSO (vehicle control), RAS inhibitor, or JAK inhibitor for 3 hrs. Cells were fixed in 4% PFA in a 96-well plate for 15 min during which the plate was centrifuged at 400×g. Cells were permeabilized with methanol at −20 °C for 5 min. After blocking with 5% FBS, primary antibodies were incubated over night at 4 °C. On the next day, rabbit and mouse probes from the Duolink In Situ Orange kit (Cat.#DUO92102; Sigma-Aldrich) or the Duolink flowPLA Orange kit (Cat.#DUO94003; Sigma-Aldrich) were used and the PLA was performed according to the manufacturer’s protocol.

Operetta CLS high-content screening microscope (PerkinElmer, Waltham, US) were used to detect and count the fluorescent PLA spots in at least 600 single cells for each well and condition. Spot distribution histograms and non-linear Gaussian fitting curves were plotted in Prism v8.1 (GraphPad Software, San Diego, US).

### Principle component analysis (PCA)

ELISA and quantified WB data for all 38 samples successfully analyzed for this study were fed into the multidimensional vectorial data visualization software ViDaExpert v1.2(23). The 13 variables included for the PCA calculations were: pan-RAS, JAK2, STAT5, MEK1/2, ERK1/2, and rpS6 activity levels in absence or presence of TSLP as well as CRLF2 protein expression. Eigenvector-based multivariate analysis was performed and the contributions of the principal components 1-3 (PC1-3) of each original variable were calculated (Supplementary-Fig.S5A). Data was transformed to a new 3D coordinate system using the projection of PC1-3. For the *k*-means clustering of the patient samples, *k* was set to 4 after identifying the smallest, significantly different class-class deviation for *k* (Supplementary-Fig.S5B). The resulting coordinate system was loaded into Adobe Illustrator software for presentation. The heatmap was generated in R-software 3.6.0 (The R Foundation, Vienna, AT) by performing unsupervised hierarchical Ward’s clustering algorithm on the DS-ALL presentation samples (using the same variables of the PCA), with correlation coefficient as distance metric.

### Statistical analysis

For all multiple comparison analysis, one-way ANOVA and post-hoc Bonferroni calculations were performed in Prism v8.1. When only two samples were compared, a two-tailed, unpaired student t-test was performed. For each series of experiments that are not independent, an additional Holm-Bonferroni correction was carried out using an Excel-macro(24) to adjust the *P*-values for sequential multiple comparison. Fisher’s exact test was calculated online(25).

Kaplan–Meier survival estimator plots and multivariate analysis using Cox proportional-hazards model were calculated in R-software 3.6.0, for details on the used factors see the respective figure legends. Variable/category candidates stated in the respective figures were included because they were either the respective analyzed protein/mutation/activity, or are known general ALL prognostic markers.

Standard protocols for the sequencing and transduction are available in Supplementary methods.

## Results

### CRLF2 and JAK2mut co-expression is sufficient to activate RAS in Ba/F3 cells

We hypothesized that increased CRLF2-level in combination with a mutation in JAK2 pathway genes could be sufficient to activate wild-type (wt) RAS protein in the absence of RAS mutations, as a mechanism to explain the mutual exclusion of JAK2 and RAS/MAPK mutations in DS-ALL. The level of RAS activity is generally assessed using a pull-down assay whereby the (activated) RAS-GTP is captured by virtue of its high affinity to RAS-binding-domain (RBD) of RAF proteins. In order to observe the effects of elevated CRLF2 signaling on the activation of RAS, we stably integrated a human *CRLF2* overexpression construct(26) into the mouse pre-B-cell line Ba/F3. This alteration did not increase the level of pulled-down RAS-GTP (Fig.1A) and neither did the stable overexpression of hJAK2_R683G_(26), the most prevalent specific activating JAK2 mutation in DS-ALL and Ph-like-ALL. However, when both of these alterations were combined, eight fold higher RAS-GTP level was measured, in the absence of cytokines (Fig.1B, post-hoc Bonferroni p-values are listed in Supplementary-Tab.S2). Growth of Ba/F3 cells depends on IL-3 (Supplementary-Fig.S1A), which induces wtJAK2 phosphorylation(27), and interestingly we found that it also activates RAS (Supplementary-Fig.S1B). The cells with combined CRLF2 and JAK2_R683G_ overexpression were the only ones in this series that grew in a cytokine-independent manner (Supplementary-Fig.S1E), as also previously observed(26). This proves that increased CRLF2-expression together with activated JAK2 is sufficient to activate wtRAS, and this coincides with the transition to cytokine-independent growth.

**Fig. 1:**
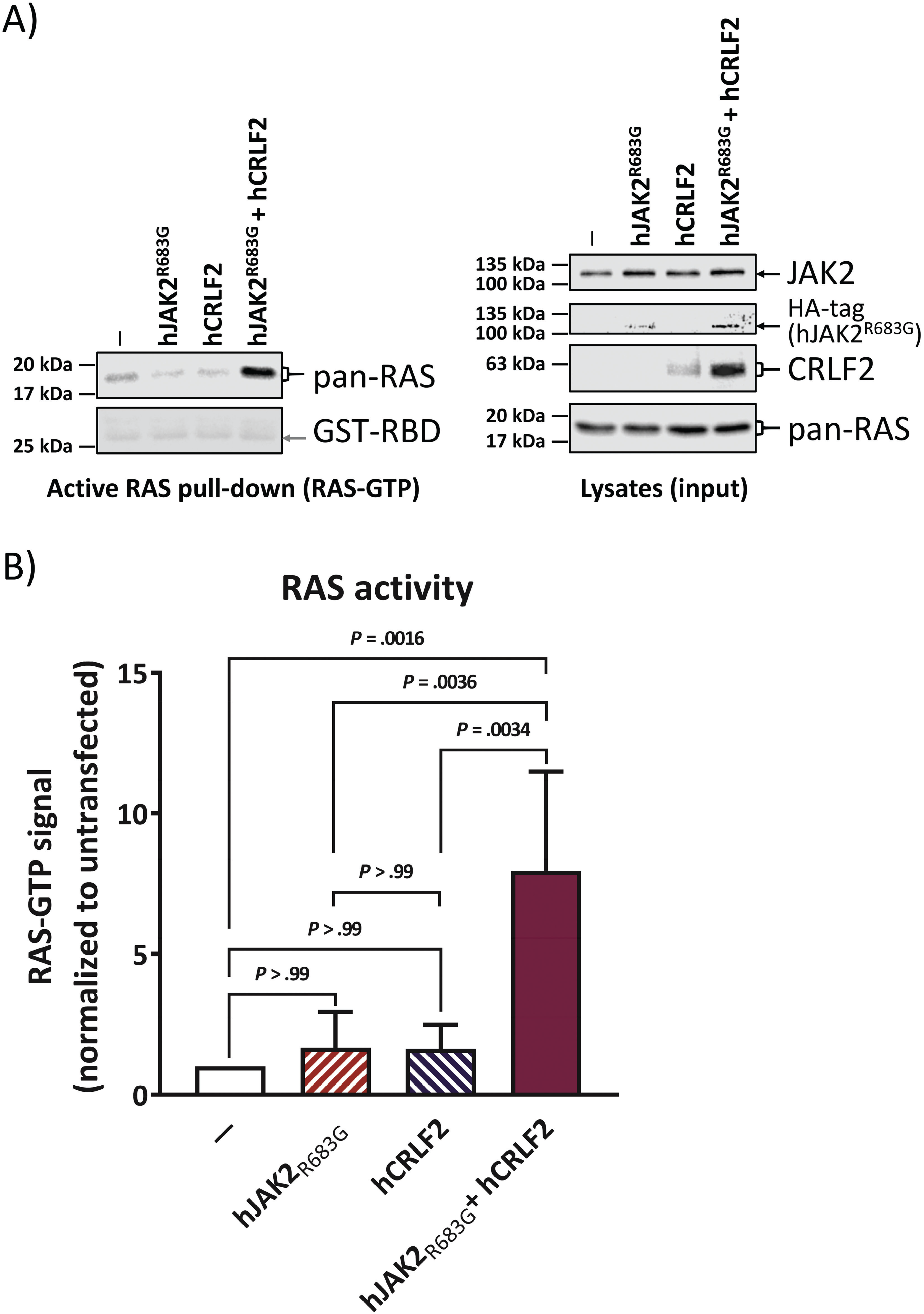
Combination of CRLF2 overexpression and constitutively active JAK2 is sufficient for wt RAS activation. Western blot analysis of the murine pro B cell line Ba/F3. Cells were stably transfected with human JAK2_R683G_ and/or human CRLF2 (see Supplementary-Fig.S1C and Supplementary-Fig.S1D) and cultured in IL-3-containing medium. All cells were then starved from IL-3 and cells were lysed. Each cell lysate was split up for analysis in RAS-GTP pull-down assay and for total proteins. An SDS-PAGE followed by Western blotting was performed. (A) Left-hand side blot shows the RAS-GTP (activated RAS) pull-down while the right-hand side blots show whole cell lysates of the same samples. Antibody-targets are labeled on the right side of each image with black arrows marking the respective protein band; the antibody against HA-tag shows the expression of the human JAK2 construct. The experiment was repeated 4 times independently. (B) Quantification of (A) for active RAS (RAS-GTP) normalized to its level in untransfected cells. Error bars are SD and *P*-values were determined in one-way ANOVA and post-hoc Bonferroni multiple comparison.

### TSLP-inducible RAS activity in absence of RAS mutations is a feature of human CRLF2 rearranged B-ALL

In order to prove the observations from Fig.1 in human ALL cells, we selected a B-ALL cell line that harbors identical changes as our double-transfected model line in Fig.1. The B cell precursor leukemia cell line MUTZ-5 from a relapsed Philadelphia-like B-ALL patient features a *CRLF2*-translocation leading to wt CRLF2 overexpression, as well as the *JAK2*_R683G_ mutation, and the absence of mutations in any RAS-MAPK pathway genes(28). The absence of *RAS* mutations in the MUTZ-5 cells grown in our cultures was confirmed by performing standard Sanger DNA-sequencing of PCR-amplicons from genomic DNA, encompassing all exons of *KRAS*, *NRAS* and *HRAS* genes (Supplementary-Tab.S3). We detected the presence of activated RAS in these cells by RAF-RBD pull-down of RAS-GTP (Fig.2A), which was tripled upon a 10 min induction with the CRLF2-ligand TSLP. Similar results were reproduced using an ELISA-based RAS-pull-down (Fig.2B). The direct binding of activated RAS and bRAF proteins in these cells was further validated via PLA (Supplementary-Fig.S2B). Both immediate upstream (PTPN11) and immediate downstream (MEK1/2, bRAF) components of the RAS/MAPK pathway were also induced by TSLP induction (Fig.2A, Fig.2B). Both KRAS and NRAS, but not HRAS, isoforms showed increased activity after TSLP-induction (Fig.2C). Interestingly, the genes for the same two isoforms (*KRAS* and *NRAS*), but not *HRAS*, acquire mutations in B-ALL(12, 20). Therefore, we conclude that the RAS isoform activity pattern of TSLP-inducibility in wtRAS leukemia cells matches the isoforms that acquire mutations in RAS-mutated leukemia cases.

**Fig. 2:**
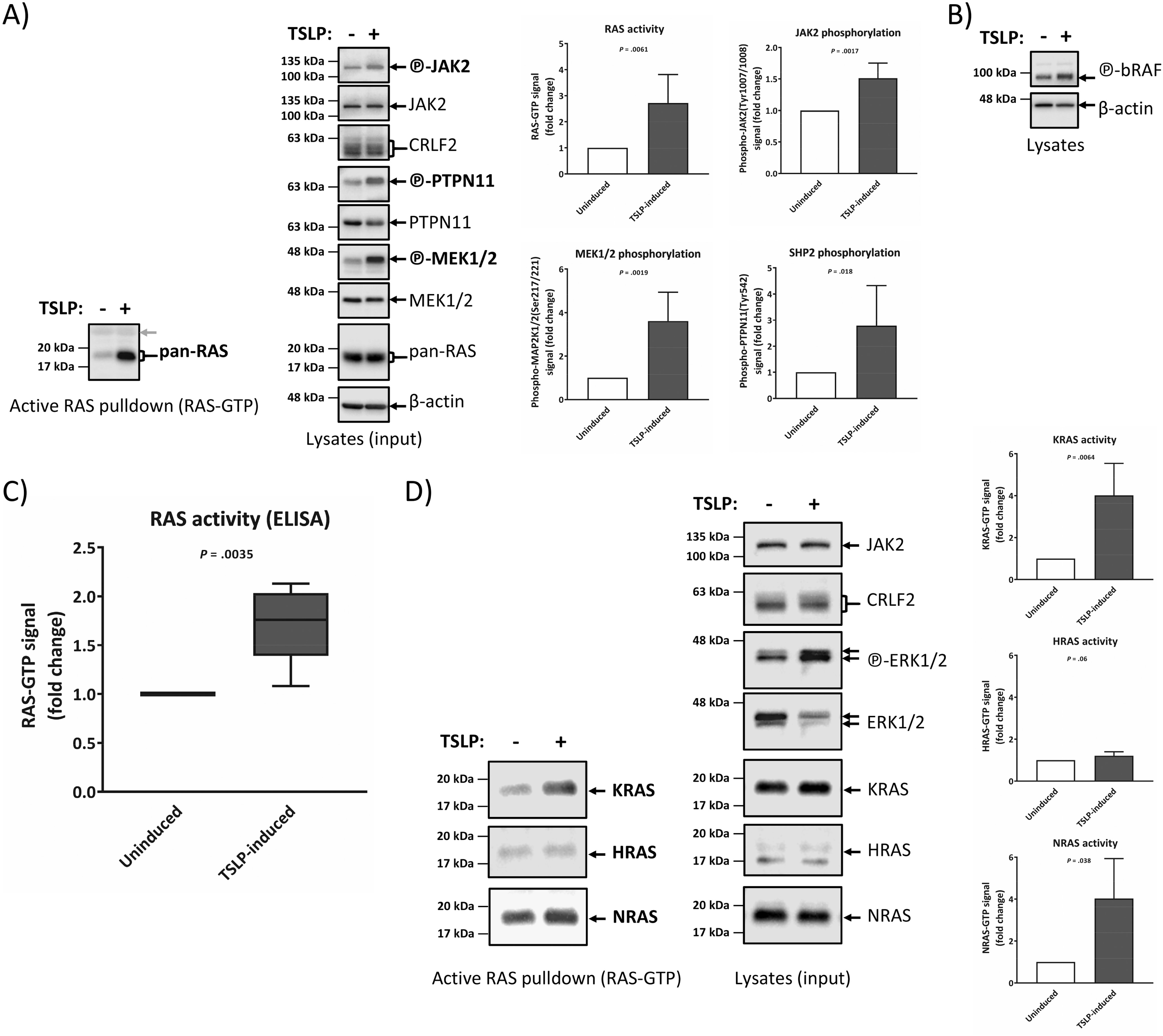
Human Ph-like B-ALL (spontaneous CRLF2-rearrangment and JAK2R683G mutation) cells activate wildtype RAS and RAS-interacting proteins upon TSLP induction. MUTZ-5 cells (Human Ph-like B-ALL cells bearing CRLF2-rearranged and spontaneous JAK2R683G mutation) were stimulated with 20 ng/mL human TSLP (maximal effective TSLP-concentration, Supplementary-Fig.S3B) for 10 min before cell lysis. Each cell lysate was split up for analysis in RAS-GTP pull-down assay and for total protein signal. (A) RAS-GTP pull-down and lysate samples were loaded on separate gels. An SDS-PAGE followed by Western blotting was performed. To assess the total protein and phosphorylated protein amounts on the same PVDF-membrane, each membrane part was stripped and reprobed with new antibodies. RAS-GTP pull-down samples are on the left side while the right-hand side blots show whole cell lysates of the same samples. Antibody-targets are labeled on the right side of each image with black arrows indicating the respective protein band. The grey arrow shows the unspecific signal of the GST-RAS binding domain (RBD) used in the active RAS pull-down assay acting as a loading control. The experiment was repeated 5 times independently and the graphs show the quantification for active RAS (RAS-GTP), phosphorylated MEK1/2 (phospho-MEK1/2), JAK2 (phospho-JAK2), and PTPN11 (phosho-PTPN11). Beta-actin and total protein signals were used as a loading control to normalize samples. (B) A blot separate from (A) demonstrates the TSLP-inducibility of RAS-effector bRAF. (C) RAS-GTP quantification of 5 independent ELISA experiments in which RAS activity of TSLP-induced MUTZ-5 cells was measured using a different, ELISA-specific active RAS pull-down assay. (D) TSLP-induced MUTZ-5 cells were probed for the presence of activated isoforms KRAS-GTP, HRAS-GTP, or NRAS-GTP (blots on the left). The blots on the right show the total expression of the respective RAS proteins and the graphs show the signal fold change over uninduced MUTZ-5 cells for KRAS-GTP, HRAS-GTP and NRAS-GTP of 4 independent experiments. All error bars are SD; *P*-values were calculated using Student’s T-test and are adjusted with a Bonferroni-correction for sequential multiple comparison.

### DS-ALL patients differ in the level of activity and inducibility of RAS, independently of RAS mutations

The MUTZ-5 ALL cells used in the analysis so far share the increased CRLF2 expression and mutated JAK2 with approximately a third of DS-ALL patients(20), which also have no mutations in RAS genes. We therefore analyzed primary cells from presentation samples (at primary diagnosis) of DS-ALL in RAS pull-down WB and ELISA assay measurements (+/− TSLP stimulation). The analyses were performed blinded to the mutation profile of the patient material and distinct DS-ALL patient profiles for RAS-activity and TSLP-inducibility of RAS were observed in WB (Fig.3A) and confirmed by ELISA (Fig.3B). As we see examples of RAS-mutated, wtRAS, or JAK2-mutated DS-ALL in each of the profile types (Fig.3C), with exception of low-RAS and non-inducible type, we can conclude that activity levels and TSLP-inducibility of RAS cannot be predicted on the basis of DNA-sequencing (acquired mutations) patterns.

**Fig. 3:**
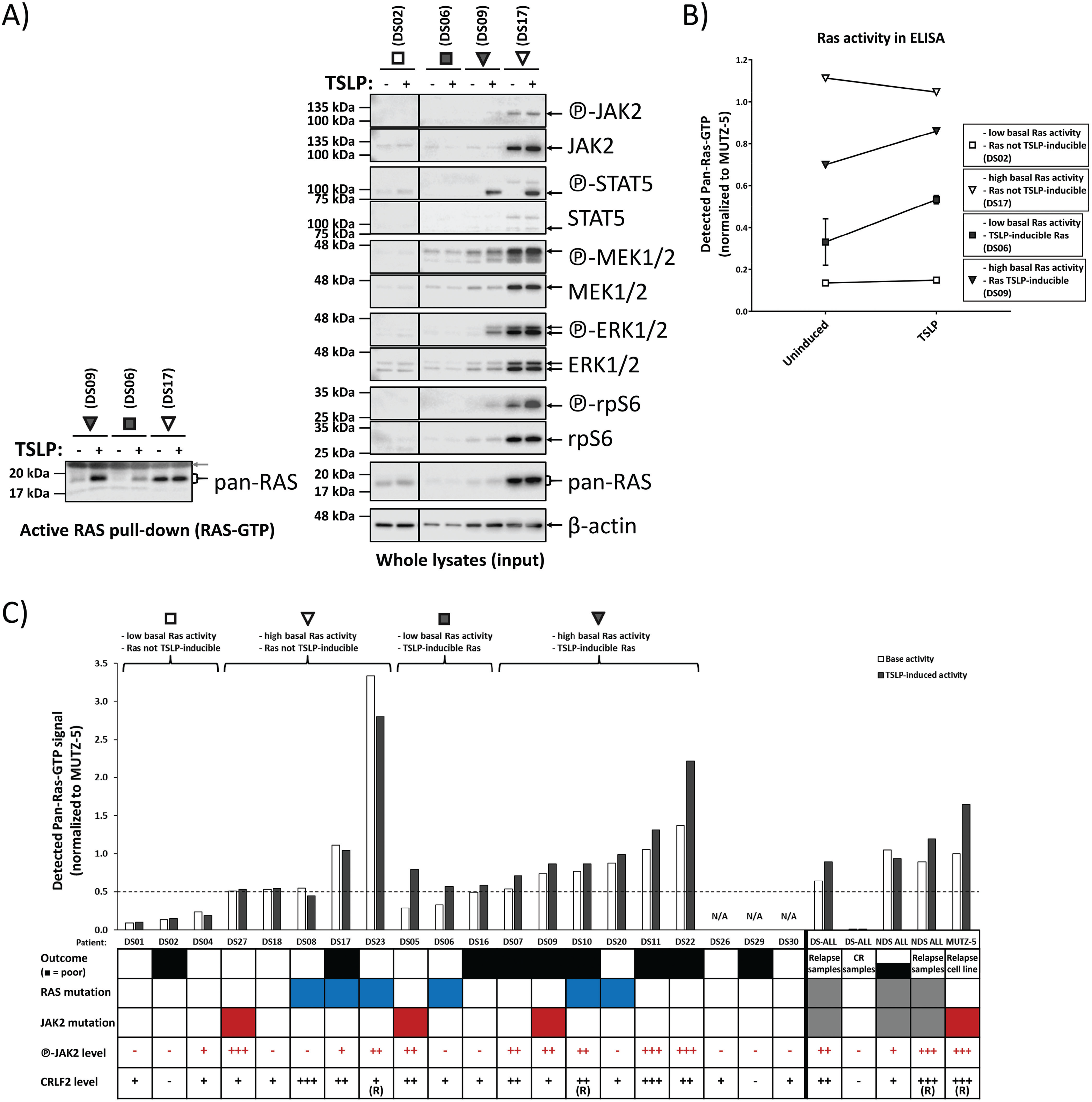
70% of primary bone marrow presentation samples of Down syndrome ALL patients show activated and/or TSLP-inducible RAS, regardless of mutations status. Primary presentation samples of DS-ALL patients were cultured for 2 days (see Supplementary-Fig.S4A legend for details) and then induced for 10 min with 20 ng/mL TSLP (or not induced) in serum-reduced medium. Lysates were analyzed for RAS activity in WB pull-down (A) or ELISA (B) using an ELISA-specific RAS pull-down assay. (A) Each lysate was split up for analysis in Western blot RAS-GTP pull-down assay and for total protein signal. RAS-GTP pull-down (left) and lysate samples (right) were loaded on separate gels. An SDS-PAGE followed by Western blotting was performed. To assess the total protein and phosphorylated protein amounts on the same PVDF-membrane, membranes were stripped and reprobed with new antibodies. Antibody-targets are labeled on the right side of each image with black arrows indicating the respective protein band; the grey arrow shows the loading of the GST-RBD in the pull-down assay. (B) The RAS activity pattern in the patient samples from (A) was confirmed via ELISA measurement of RAS-activity in aliquots that were independently thawed and processed as described above The line graph illustrates the four main patterns observed for RAS activity in primary ALL patient samples. (C) Shows an overview of the ELISA-measured RAS activity for the DS-ALL cohort at diagnosis (not enough cell material was available for DS26, DS29 and DS30). The RAS-GTP pull-down for ELISA was performed on lysates from cells at minimum 75% viability at a 100 ng/μL total protein concentration. In parallel, uninduced MUTZ-5 cells were subjected to the same treatment as the primary patient cells and were used to normalize all RAS activities. Brackets on top indicate the groups of the four RAS activity patterns presented in (A, B). For visualization purposes only in this graph, basal RAS activity over 50% of MUTZ-5 basal RAS activity was grouped as high RAS activity while an increase by at least 10% RAS-GTP in TSLP-stimulated samples over uninduced samples in ELISA was classed as TSLP-inducible RAS. For visualization purposes only in this graph, phosphorylation levels measured in WB for JAK2 were categorized as –(negative) = 0.00-0.05; + = 0.05-0.50; ++ = 0.50-1.00; +++ =1.00-2.00, and CRLF2 protein levels were categorized as –(negative) = 0.00-0.05; + = 0.05-0.20; ++ = 0.20-0.50; +++ =0.50-1.50. None of the arbitrary threshold groupings defined above were used in any of the PCA or clustering analysis shown later (Fig.3, Supplementary-FigS5). Known CRLF2-rearrangements are marked (R). All values are normalized to those measured for uninduced MUTZ-5 cells.). Outcome of leukemia is given (white = good outcome, black = poor outcome), and the presence of *RAS* mutations (blue) or *JAK2* mutations (red) are specified (grey means unsequenced samples). The groups at the right end of the bar graph (separated by the black bar) show average RAS activities for patient/sample groups other than DS-ALL-diagnosis: Non-DS (NDS) at presentation, DS complete remission (CR) and DS/NDS at relapse. For an overview of the complete Western blot data and the quantified activities and protein expression of STAT5, JAK2, MEK1/2, ERK1/2 and rpS6 of all individual samples, see Supplementary-Fig.S4.

The most important conclusion of this analysis is that in 14/20 (70%) of primary DS-ALL samples analysed, 8 of which had no RAS mutations, but 75% of those had either mutated or hyperphosphorylated JAK2 (Fig.3C). This means that either the RAS mutation, or the combination of high CRLF2 and hyperphosphorylated JAK2 (including mutated JAK2) can explain the mechanism for high RAS activity in 12/14 (86%) of DS-ALL with high RAS activity. Importantly, not a single wtRAS case with either mutated or hyperphosphorylated JAK2 was seen that lacked activated RAS protein (Fig.3C).

### RAS activity and its TSLP inducibility correlate with outcome in DS-ALL patients

Data from primary cells analysis from n=20 presentation samples of DS-ALL for the RAS/MAPK, PI3K/mTOR, and JAK/STAT pathway activity profiles using WB (Supplementary-Fig.S4), as well as ELISA for activated RAS-pull-down (Fig.3C) were integrated with the similar data we obtained using n=7 DS-ALL relapse and n=4 DS-ALL remission samples, as well as n=4 non-DS ALL presentation samples and n=2 non-DS relapse samples. We performed a PCA using all of these integrated data on N=37 samples from n=31 individual patients, in parallel to the same readouts from the MUTZ-5 Ph-like ALL reference cell line, and the PCA result was mapped onto a coordinate system (Fig.4A) using the three principle components (PC1-3, Supplementary-Fig.S5A). Unsupervised *k*-means clustering grouped ALL samples into Clusters 1-4 (Fig.4A). This analysis grouped almost all presentation and remission samples of 9-year event-free survival patients (good outcome) together into Cluster 1 (green symbols). In contrast, out of 15 samples grouped into Cluster 2 (red symbols), 13 samples (87%) were from patients with death or subsequent relapse as outcome. Clusters 3 and 4, further along the PC1-axis, consisted of a small number of exclusively relapse samples. Using an independent mathematical approach, unsupervised hierarchical clustering of the 20 DS-ALL presentation samples (Supplementary-Fig.S5D) grouped 90% of the samples into the same groups as the PCA-mapping. The clustering revealed that presentation samples from Cluster 1 correlated with good outcome for DS-ALL patients while DS-ALL patients grouped into Cluster 2 showed a significantly increased risk of relapse (Fig.4B,C). The PCA-derived protein activity score was independently predictive of outcome (*P*=.041) (Fig.4C) when analyzed by a multivariate Cox regression model together with CRLF2 protein-expression, NCI-risk and JAK2-mutation status (or RAS-mutation status, not shown).

**Fig. 4:**
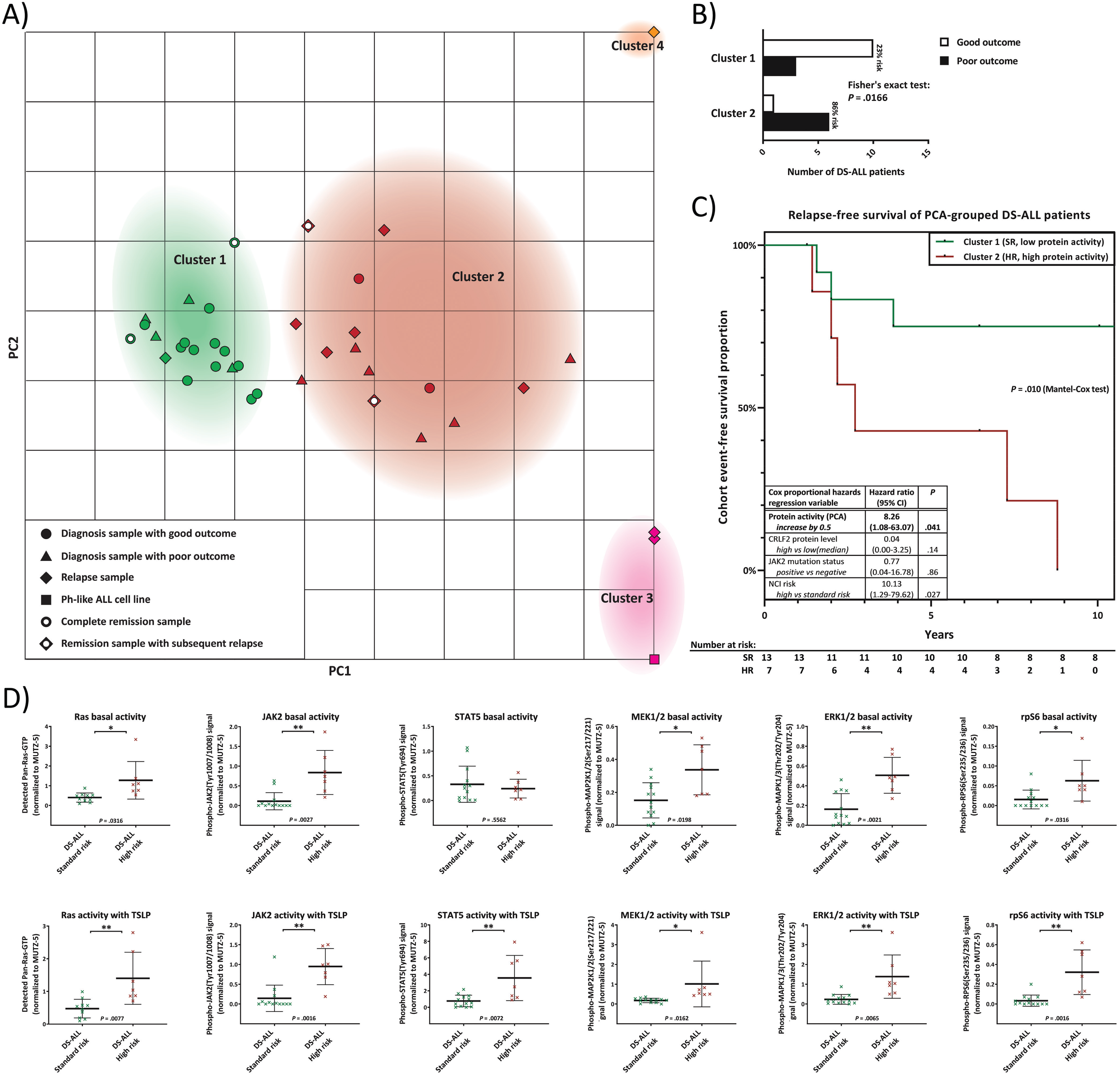
Sub-stratification of DS-ALL patients based on primary cells at diagnosis: patterns of RAS activation, TSLP-inducibility and downstream signaling in relation to standard therapy outcomes. (A) A principal component analysis (PCA) was performed on the quantified data of Fig.3 (data was given as continuous variables; no cutoffs or pre-grouping were used) for the DS-ALL cohort at diagnosis, and (where available) at remission and relapse, as well as primary presentation and relapse samples from Non-DS ALL patients. Top view of the PCA mapping for all 6 measured protein activities (basal and TSLP-induced) as well as CRLF2 protein expression of all analyzed samples along the calculated principle components (for PCA calculated results see Supplementary-Fig.S5A). *K*-means clustering (with *k* set to 4 to achieve minimal class-class deviation, Supplementary-Fig.S5B) grouped samples into clusters 1 to 4 (listed in Supplementary-Fig.S5C). Sample data points are colored according to the unsupervised clustering. (B) PCA Clusters 1 and 2 contain all samples of the DS-ALL diagnosis cohort and were analyzed according to their outcome: A Fisher’s exact test determined the *P*-value between the number of good and poor outcomes between the two clusters (bar graph). (C) Kaplan–Meier curves of cluster 1 (SR = standard risk) and cluster 2 (HR = high risk) DS-ALL patients. Table show a Cox proportional-hazards model for protein activity score (PCA-derived principle component from all quantified protein activities at basal and TSLP-induced level) together with CRLF2-protein expression level (for CRLF2+ samples), NCI risk groups (SR: age at diagnosis 1-10 yr and WBC < 50.000/µL; HR = children age > 10 yr and/or WBC > 50.000/µL; or unknown), and presence of activating JAK2 mutations. Reverse Kaplan-Meier median follow-up for N=20 DS-ALL was 18.4 years. Patient numbers at risk for each year are given in the table below the survival curve. (D) The means of all analyzed basal or TSLP-induced protein activities are compared between the SR group (DS-ALL patients in PCA cluster 1) and the HR group (DS-ALL patients in PCA cluster 2). All error bars are SD; *P*-values were calculated using Student’s T-test and are adjusted with a Bonferroni-correction for sequential multiple comparison. For visibility, significant *P*-values were additionally highlighted with * (*P* < .05) or ** (*P* < .01).

We then restricted the further analysis only to DS-ALL primary presentation samples, and quantitatively compared those that PCA grouped into Cluster 1 (PCA-predicted standard-risk (SR)) to those in Cluster 2 (PCA-predicted high-risk (HR)), for the basal activities (Fig.4D top row)and TSLP-induced activities (Fig.4D bottom row) of pan-RAS, JAK2, STAT5, MEK1/2, ERK1/2 and rpS6. We observed that basal and TSLP-induced activities of JAK2, ERK1/2 and rpS6 were significantly increased in HR-DS-ALL presentation samples compared to the SR group (within PCA-Cluster 1). For these proteins, correlation between risk and protein-activity/inducibility profile for our DS-ALL cohort, resembles previously reported findings for a different group of HR-ALL, the non-DS Ph-like ALL, grouped by the presence or absence of CRLF2 rearrangements(29). Additionally, (and this has, to our knowledge, never been demonstrated for any ALL before), we also observed a significant increase in basal and TSLP-induced activity of both MEK1/2 and RAS in the HR-DS-ALL group, compared to the SR group. We also looked at protein expression levels and found RAS and rpS6 levels to correlate with the high-risk DS-ALL group (Supplementary-Fig.S6A). This provided the rationale to look for differences in the larger non-DS ALL cohort (346 children treated on the Malaysia-Singapore ALL protocols), for which unfortunately no protein material was available. The available material only permitted an analysis of mRNA transcript levels of the same genes within the whole transcriptome RNA-seq data for this cohort (N=346). Interestingly, in the subgroup of this cohort with CRLF2-expressing, high-risk non-DS ALL (n=91; median follow-up = 6.95 years), KRAS mRNA-expression was independently predictive of outcome (*P*=.030) when included in a multivariate analysis using Cox proportional hazards regression model together with Ph-like status, RAS mutation status, sex, and NCI risk (Supplementary-Fig.S7B).

The combined data suggest that a large scale analysis on a non-DS ALL cohort is warranted for protein activation patterns of RAS, MEK1/2, and other pathway components activation readouts as this could potentially significantly inform the patient sub-stratification for outcome.

### RAS inhibitor can significantly block the growth of human B-ALL Ph-like wtRAS cells

We next examined to what extent the direct RAS activation in wtRAS leukemic cells affects the cell growth and viability. We tracked the cell count and cell viability of MUTZ-5 cells after treatment with pan-wtRAS-inhibitor and in comparison to treatments with other compounds that have been previously reported to induce dose-dependent cytotoxicity in MUTZ-5(29), some of which are currently in clinical trials for Ph-like ALL(6). After 4 days treatment with pan-wtRAS inhibitor the growth and viability of MUTZ5 cells were significantly reduced (Fig.5A,B), and this was not affected by the presence of TSLP. In comparison, the dual PI3K/mTOR inhibitor also significantly reduced the growth and viability of MUTZ-5 cells, but this inhibitory effect could be partially counteracted by the TSLP-induction (Fig.5A). Both of these compounds, at the concentrations used (chosen to achieve almost full efficacy on the respective main pathway target in WB, Fig.5C), showed a much stronger inhibitory effect on cell growth than the JAK inhibitor (Fig.5A), despite the observation that the JAK inhibitor, at the same concentration, showed the strongest inhibition of TSLP-induced phosphorylation of STAT5, MEK1/2, PTPN11, ERK1/2 and rpS6 (Fig.5C, right-hand side blots). However, neither the JAK inhibitor, nor the PI3K/mTOR inhibitor could block the wtRAS activation by TSLP (Fig.5C, left-hand side blots). For the JAK2 inhibitor this might be explained by its failure to reduce the direct interaction between RAS and PTPN11 in PLA (Supplementary-Fig.S2E). In contrast, the pan-wtRAS-inhibitor completely blocked the TSLP-induced RAS activity (Fig.5C, left-hand side blot). Moreover, the pre-inhibition of RAS-direct interacting proteins (RAF and PTPN11) also reduced TSLP-induced wtRAS boost in human Ph-ALL cells (Fig.5C, Fig.8B, left-hand side blot), concordant with the PTPN11-inhibitor blocking the direct interaction between RAS and p-PTPN11 (Fig.8C). Combined, our data suggest that TSLP-activation of RAS in the absence of RAS mutations drives B-ALL cell growth, and represents an independent drug target, in addition to the PI3K/mTOR and JAK/STAT pathway targets.

**Fig. 5:**
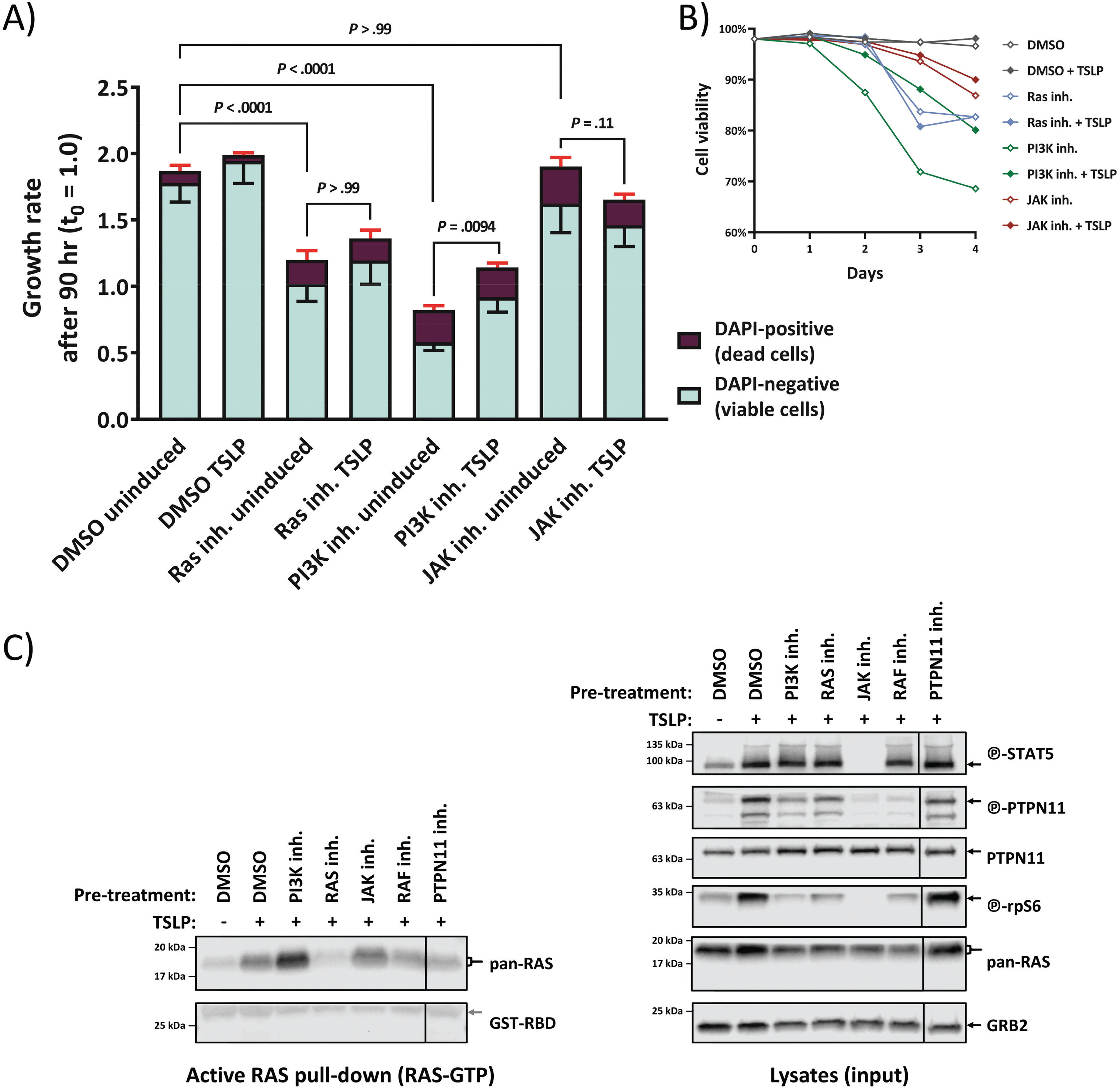
Inhibition of RAS stops wt-RAS sequence Philadelphia-like ALL cell growth in the presence of TSLP. (A) MUTZ-5 cells were seeded at 6.5×10^5^/mL density and cultured over 4 days with either 0.5% DMSO (vehicle control), 50 µM Salirasib (indirect Pan-RAS inh.), 10 µM PI-103 (PI3K/mTOR dual inh.), or 5 µM Ruxolitinib (JAK inh.), each in absence or presence of 20 ng/mL human TSLP. Cell count and viability (percentage of acridine orange-positive cells not stained by 4’,6-diamidino-2-phenylindole (DAPI) was determined in a NC-250 automated cell counter daily. The stacked-bar graph on the left side shows the growth rate after the 90 hrs timepoint, averaged from 2 independent experiments, each with triplicate wells. Red error bars are SD from the dead cell fraction while the black error bars show the SD of the viable cells. *P*-values were calculated in one-way ANOVA from the total cell growth rate and adjusted in a post-hoc Bonferroni multiple comparison. Only relevant *P*-values are shown in the graph, for a complete list see Supplementary-Tab.S2. (B) The graph shows the cell viability of the experiment in (A) over time. (C) MUTZ-5 cells were pre-treated for 2 hrs with either 0.5% DMSO (vehicle control), 10 µM PI-103 (PI3K/mTOR dual inh.), 50 µM Salirasib (indirect Pan-RAS inh.), 5 µM Ruxolitinib (JAK inh.), 50 µM Vemurafenib (Pan-Raf inh.), or 25 µM II-B08 (PTPN11 inh.), and then stimulated with 20 ng/mL human TSLP for 10 min followed by cell lysis. Each lysate sample was split up for analysis in RAS-GTP pull-down assay and for total protein signal. RAS-GTP pull-down (left) and lysate samples (right) were loaded on separate gels. An SDS-PAGE followed by Western blotting was performed. To assess the total protein and phosphorylated protein amounts on the same PVDF-membrane, membranes were stripped and reprobed with new antibodies. Antibody-targets are labeled on the right side of each image with black arrows indicating the respective protein band.

### Potent response to wtRAS-inhibitor in vitro is a distinguishing feature for poor outcome sub-cohorts of primary DS-ALL patient biopsies

We used primary surplus clinical material in Fig.4 from n=31 patients. Out of these, we had enough primary diagnosis material for 13 patients (before any therapy) to measure the effects of RAS, PI3K, or JAK inhibitors on individual pathway activation status in the presence of TSLP. The efficacy of RAS inhibition on intracellular protein activity (expressed as panRAS activity ratio between inhibitor and vehicle treated samples) for primary presentation samples showed a significant difference (*P*=.021 by Fisher’s exact test) between the good outcome (n=7) and poor outcome DS-ALL groups (n=6) (Fig.6A). Also, samples in which RAS can be further active by TSLP were more sensitive to RAS inhibitor treatment (Supplementary-Fig.S8). For all poor outcome DS-ALL primary presentation samples, inhibitions of individual pathway effector activities via the RAS, PI3K, or JAK inhibitors was visualized as inverted bar graph ranging from 0% (no inhibition) to 100% (complete inhibition) (Fig.6B). As shown by red asterisks, PI3K inhibitor significantly inhibited rpS6 phosphorylation, whereas JAK inhibitor significantly inhibited ERK, rpS6, and STAT5 phosphorylation. Notably, these inhibitors did not have any significant effect on RAS activity (absence of red asterisks), reproducing the result obtained for the MUTZ-5 Ph-like ALL cell line (Fig.5A). Only the RAS inhibitor was able to significantly block RAS activation in poor outcome DS-ALLs (Fig.6B), in addition to blocking rpS6 phosphorylation, as likewise shown for the MUTZ-5 cells (Fig.7B and Fig.5B). This suggests that only RAS-inhibitor action is capable of efficiently blocking RAS activation in cells from both Ph-like/non-DS and DS-ALL poor outcome patient samples at the point of first clinical presentation, irrespective of the presence of RAS mutation. In contrast, JAK and PI3K inhibitor treatments alone did not significantly impact RAS activity in these samples (Fig.6B). One example highlights the need for an even deeper understanding of the driving mechanisms, and the need to block wtRAS activation as part of the combinatorial treatment design. Exome sequencing found for DS-ALL patient DS09 at presentation a JAK2 mutation (and no RAS mutation). These blast cells were highly TSLP-inducible for RAS activity, which could be blocked very efficiently using RAS inhibitor, but not using JAK or PI3K inhibitors in vitro. The same patient relapsed two years after standard chemotherapy regimen, and the relapse sample contained no JAK2 mutations anymore, but the main blast had gained an NRAS mutation (sequenced as sample 4-1036101-T2(20)). Interestingly, this relapse sample with constitutively activated NRAS shows no rpS6 activity any longer (with or without TSLP-induction) and TSLP could only induce moderate levels of STAT5 activity compared to the presentation sample (Supplementary Fig.S4A).

**Fig. 6:**
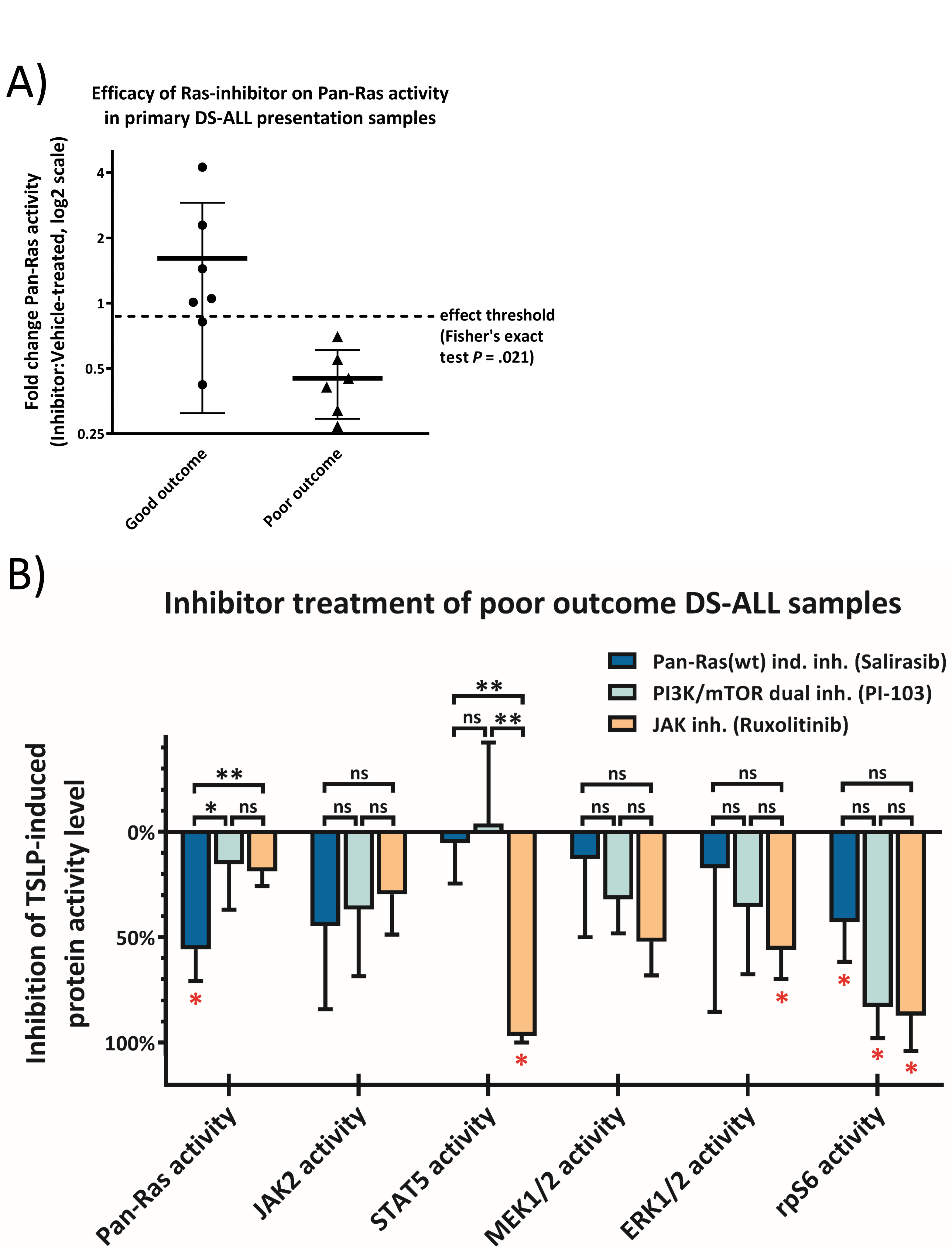
RAS inhibitor blocks RAS activity with far greater efficiency in primary, poor outcome DS-ALL patient samples, prior to relapse. Primary presentation samples of DS-ALL patients were cultured for 2 days (see Supplementary-Fig.S4A legend for details). Samples with sufficient cell count were treated with either 0.5% DMSO (vehicle control), 50 µM Salirasib (indirect Pan-RAS inh.), 10 µM PI-103 (PI3K/mTOR dual inh.), or 5 µM Ruxolitinib (JAK inh.) for 3 hrs after which the cells were induced for 10 min with 20 ng/mL TSLP in serum-reduced medium. Cells were lysed and each lysate was split up for analysis in RAS-GTP pull-down assay and for whole lysate protein signal via Western blot (Supplementary-Fig.S4A). Protein activities of all inhibitor-tested TSLP-induced samples were normalized to the activity level of the respective vehicle-treated TSLP-induced patient samples. (A) shows the efficacy of the RAS inhibitor on ELISA-measured RAS activity in DS-ALL patients with good outcome compared to those with poor outcome. If inhibitor treatment reduced the RAS activity by over 10% compared to vehicle-control (dashed line in plot), the sample was tallied as successful RAS blocking. A Fisher’s exact test was performed between the groups. The tested good outcome group (N=7) contained 3 *RAS* mutations and 1 *JAK2* mutation while none of the 6 poor outcome samples featured any *RAS* mutations (1 sample contained a *JAK2*-mutation). (B) Waterfall plot shows the efficacy (0% = no effect, 100% = full block of the respective protein activation in presence of TSLP) of all three inhibitors tested on primary presentation samples from poor outcome DS-ALL patients. Error bars are SD and *P*-values of black asterisks were determined in one-way ANOVA with post-hoc Bonferroni multiple comparison. Black asterisks (*: *P* < .05; **: *P* < .01) indicate significance comparing the three inhibitors (ns = not significant), while the red asterisks indicate if the inhibitor on average significantly reduced the respective protein activity in the primary presentation samples (absence of red asterisk = not significant; see Supplementary-Tab.S2 for a full list of Bonferroni-corrected p-values). *P*-values of red asterisks were calculated using Student’s T-test and are adjusted with a Bonferroni-correction for sequential multiple comparison.

**Fig. 7:**
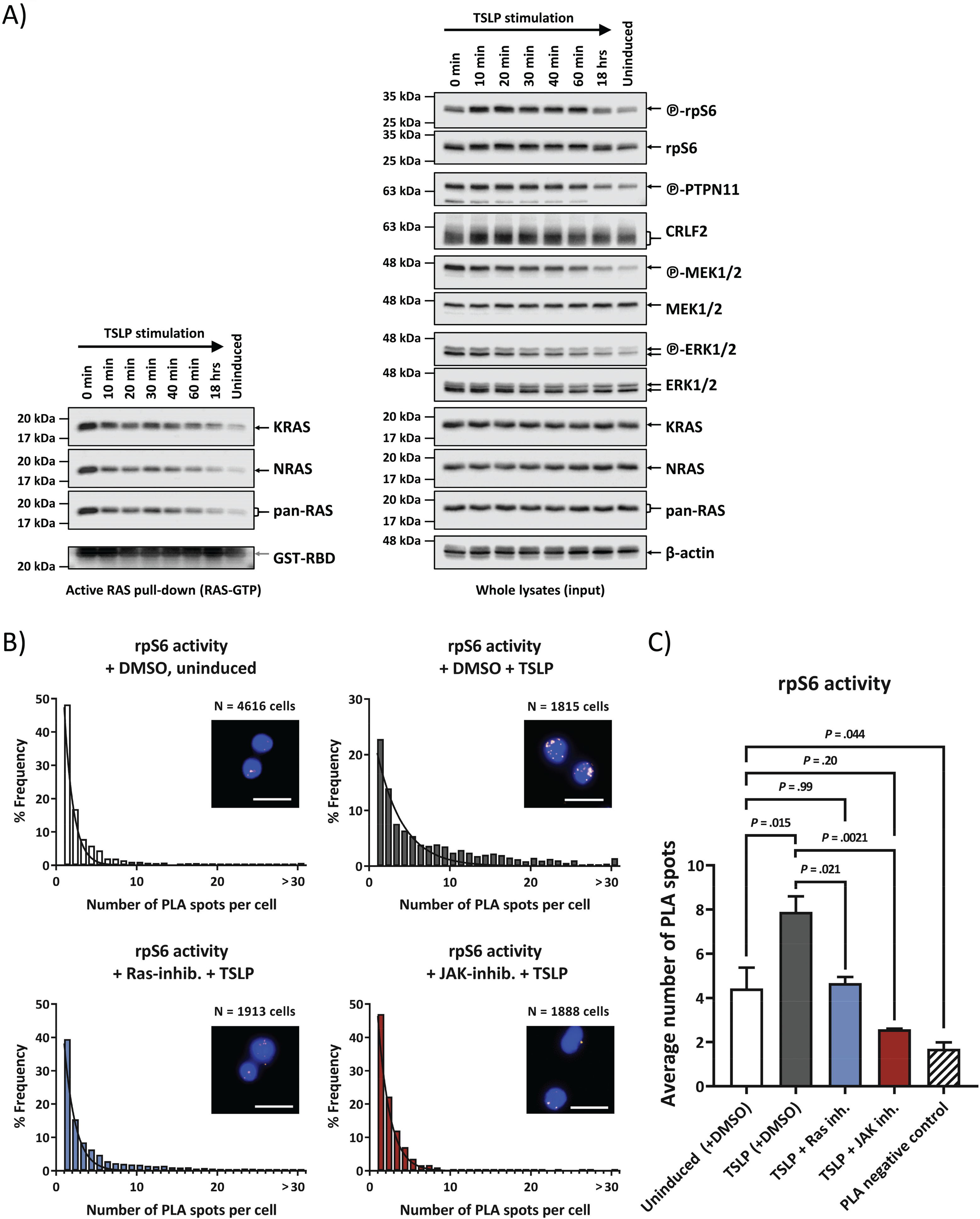
Direct wtRAS activation can precede PI3K/mTOR pathway activation, and the resulting PI3K downstream signaling activity can be blocked by RAS inhibitor. (A) Effect of TSLP induction over time. MUTZ-5 cells were incubated with 20 ng/mL human TSLP at 37 °C and 5% CO_2_ for the indicated time points (0 min to 18 hrs) before cell lysis. Due to the centrifugation step of the suspension cells the TSLP is able to act for 5 min before lysis at timepoint 0. Each cell lysate was split up for analysis in RAS-GTP pull-down assay and for total protein signal. RAS-GTP pull-down and lysate samples were loaded on separate gels. An SDS-PAGE followed by Western blotting was performed. To assess the total protein and phosphorylated protein amounts on the same PVDF-membrane, membranes were stripped and reprobed with new antibodies. RAS-GTP pull-down elutions are on the left side while the right-hand side blots show whole cell lysates of the same samples. Antibody-targets are labeled on the right side of each image with black arrows indicating the respective protein band. (B) Activation of PI3K/mTOR downstream target rpS6 protein was monitored via PLA in high-throughput microscopy. MUTZ-5 cells were either not induced or induced with 20 ng/mL TSLP for 10 min. Where indicated, cells were pre-treated for 3 hrs with either DMSO (vehicle control), RAS inhibitor, or JAK inhibitor. Cells were fixed and permeabilized in a 96 well plate. After blocking, antibodies against phosphorylated rpS6 and total rpS6 were used in conjunction with PLA rabbit and mouse probes to allow specific readout of rpS6 activation in single cells in a high-throughput manner. Histograms show the distribution for a single experiment of the number of PLA spots in cells with at least 1 PLA spot (assay control is only shown in the bar graph). A minimum of 600 cells were analyzed per sample. Non-linear Gaussian fitting curves were plotted. Fluorescent microscope images show examples of PLA spots in MUTZ-5 cells for the respective treatment; white scale bars are 20 µm long. (C) The bar graph summarizes the average PLA spot counts of 3 independent experiments. Error bars are SD and *P*-values were determined in one-way ANOVA and post-hoc Bonferroni multiple comparison.

Our data on primary patient material suggest the compulsory activation of RAS whenever elevated CRLF2 is present in combination with either mutated or otherwise activated JAK2. This would eliminate the selective advantage gained by a RAS mutation, explaining the mutual exclusion, however the underlying molecular mechanism remains to be explained. We therefore sought to further characterize the molecular mechanism behind the wtRAS activation in these leukemic cells.

### TSLP activates RAS directly and independently of PI3K/mTOR pathway activation

The use of inhibitors on patient samples (Fig.6B) suggested RAS activation to be independent from blocking of PI3K or JAK pathways. TSLP induction in high CRLF2-expressing and JAK2-mutated B-ALL is known to activate STAT5 and PI3K/mTOR pathways(29), and this insight is exploited in innovative new therapeutic approaches that are currently clinically trialed(6). We therefore first confirmed that our experimental system can reproduce these same results (Supplementary-Fig.S3A).

Downstream effectors of the activated PI3K/mTOR pathway have been shown in some situations to cross-activate the downstream effectors of RAS-MAPK cascade, and vice versa(30, 31). However, we observed that immediately upon addition of TSLP (0 min timepoint, Fig.7A) the relative levels of activated pan-RAS, KRAS, NRAS, PTPN11, MEK1/2, and ERK1/2 were all higher than at any later timepoint, while in comparison, the activity onset of the PI3K/mTOR downstream target rpS6 was delayed (Fig.7A). This makes it less likely, at least as the initial effect of TSLP, that the activation of MEK1/2 and ERK1/2 in such leukemic cells is caused by the cross-talk from the activated PI3K pathway. As PI3K can also be an effector of RAS(32), we used an alternative biochemical approach (PLA) by which we demonstrated the ability of a chemical inhibitor of RAS (Salisarib) to block the TSLP-induced rpS6-activating phosphorylation (Fig.7B,C), at a concentration lower than required to block EIF4EBP1 activity via mTOR-complex destabilization(33). PLA also detected a strong interaction between RAS and the RBD-containing PI3K-subunit p110α in these cells, which could be reduced using Rigosertib, a RAS-GTP mimetic that inhibits RAS by binding to the RBD of RAS-effectors (Supplementary-Fig.2D).

Our data therefore strongly suggest that direct, wtRAS activation can precede, and to a certain extent promote, the PI3K/mTOR pathway activation in TSLP-induced human ALL cells.

### CRLF2-signaling increases direct interaction between active PTPN11 and RAS

Active PTPN11 is known to dephosphorylate RAS to prime it for activation(34), and we found PTPN11 phosphorylation to be increased by induced CRLF-signaling (Fig.2A). Furthermore, PTPN11 is published to be in complex with JAK2 upon cytokine-induction in tumor cells(35). In order to confirm that the mechanism of activating RAS in JAK2-mutated B-ALL cells is regulated via PTPN11, we designed a PLA assay that specifically detects the direct interaction between RAS and phosphorylated PTPN11 (Fig.8A,C). Indeed, compared to the signal for two cytosolic proteins not expected to interact (PLA negative control (NC)), a strong PLA signal between RAS and p-PTPN11 was observed and this interaction almost doubled upon TSLP-induction (Fig.8A). PLA assays also detected interactions between SOS1 and GRB2 in these leukemic cells, as well as other direct interactions involved in RAS activation, which also showed response to CRLF2-activation (RAS and SOS1; GRB2 and p-PTPN11) (Supplementary-Fig.S2C). Remarkably, blocking PTPN11-activity via the PTPN11-inhibitor II-B08 reduced both endogenous, and TSLP-induced RAS activity in these cells (Fig.8B). The PTPN11-inhibitor did not reduce the phosphorylation marker on PTPN11 itself (Fig.8B), but disrupted the direct interaction between RAS and p-PTPN11, lowering it to levels below those in uninduced cells (Fig.8C). Furthermore, a cytotoxic assay showed leukemic cell growth and viability to be reduced by the PTPN11-inhibitor, similarly as with the RAS-inhibitor (Fig.8D). Taken together, these results show that the mechanism of wtRAS activation by CRLF2 signaling depends on its direct interaction with catalytically active PTPN11.

**Fig. 8:**
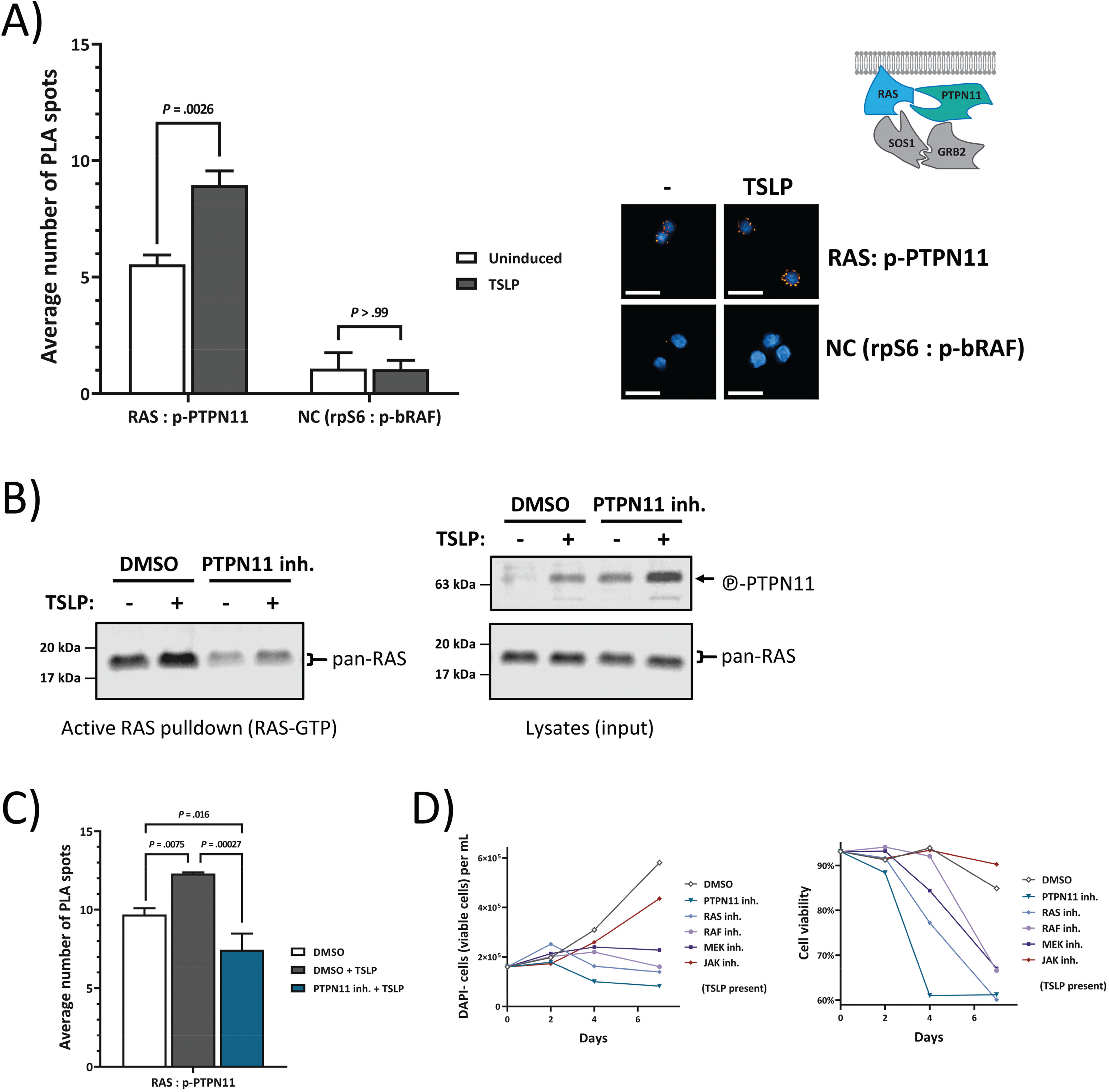
CRLF2-signaling induces direct interaction between activated PTPN11 and RAS, and PTPN11 activity is required for RAS-dependent ALL cell growth. (A) Direct interaction between RAS and phosphorylated PTPN11 (see cartoon in top right corner) was monitored via proximity ligation assay (PLA) using high-content microscopy. Serum-starved MUTZ-5 cells were either not induced or induced with 20 ng/mL TSLP for 10 min. Cells were fixed and permeabilized in a 96 well plate. After blocking, antibodies against phosphorylated PTPN11 and pan-RAS were used in conjunction with PLA rabbit and mouse probes to allow the amplification and staining of interaction-specific PLA spots. The negative control (NC) samples used antibodies for two cytosolic proteins (rpS6 and phosphorylated bRAF) that were not expected to directly interact. Fluorescent microscope images show examples of PLA spots for the respective treatment; white scale bars are 20 µm long. At least 700 cells per technical triplicate were analyzed in a high-throughput manner using Operetta automated high-content microscopy platform. The bar graph shows the averages of 3 independent experiments (each performed in triplicates). Error bars are SD and *P*-values were determined in one-way ANOVA and post-hoc Bonferroni multiple comparison. (B) MUTZ-5 cells were pre-incubated with DMSO or 25 µM II-B08 (PTPN11 inh.) for 2 hrs and then stimulated or not with 20 ng/mL human TSLP for 10 min before cell lysis. Each cell lysate was split up for analysis in RAS-GTP pull-down assay and for total protein signal. RAS-GTP pull-down and lysate samples were loaded on separate gels. An SDS-PAGE followed by Western blotting was performed. RAS-GTP pull-down samples are on the left side while the right-hand side blots show whole cell lysates of the same samples. Antibody-targets are labeled on the right side of each image with black arrows indicating the respective protein band. (C) MUTZ-5 cells were pre-incubated with DMSO or 25 µM II-B08 (PTPN11 inh.) for 2 hrs and then stimulated or not with 20 ng/mL human TSLP for 10 min before the PLA described in (A) was performed. (D) MUTZ-5 cells were seeded at 1.6×10^5^/mL density and cultured over 7 days with either 0.5% DMSO (vehicle control), 50 µM Salirasib (indirect pan-RAS inh.), 25 µM II-B08 (PTPN11 inh.), 50 μM Vemurafenib (Pan-Raf inh.), 1 μM PD0325901 (MEK1/2 inh.), or 5 µM Ruxolitinib (JAK inh.), in presence of 20 ng/mL human TSLP. Live cell count (left graph) and viability (percentage of acridine orange-positive cells not stained by 4’,6-diamidino-2-phenylindole (DAPI)) (right graph) was determined in a NC-250 automated cell counter.

## Discussion

Both DS-ALL and Ph-like ALL share CRLF2-rearrangements and various kinase-activating alterations as potential targets for individualized therapy using specific kinase-inhibitors(8, 36). This lead to the use of phosphorylation patterns of individual kinase signaling cascades as informative biomarkers for combinatorial therapy design(6, 16, 37).

In DS-ALL, recent studies of sub-clonal and single-cell evolution of changes in leukemic ALL blasts have identified signaling activators (CRFL2-rearrangements, JAK2 mutations, RAS-MAPK mutations and iAMP21) as frequent events in primary and relapsed leukemic blasts(20, 21, 38). In particular, JAK2 and RAS mutations were found to be both acquired and lost in relapse samples in a mutually exclusive manner(20, 21). This emphasizes the need for individualized combined-therapy approaches that have a better chance of preventing the selection of sub-clones driven by a different signaling. Our data show that elevation in CRLF2 levels combined with JAK2 activation are sufficient to activate wtRAS, and that TSLP has the potential to induce the wtRAS activity, independently of the PI3K/mTOR activity. This has implications on the choice of the combinatorial therapy design. Remarkably, our combined data from exome sequencing(20), and primary ALL cell protein signaling (presented in this study), suggest that up to 65-82% of DS-ALL cases have highly activated RAS, either constitutively, or upon TSLP induction, regardless of their mutation profiles. 12 of 14 cases with high RAS-activity featured either RAS mutations or high CRLF/JAK2 signaling (including JAK2-mutations). The only two samples featuring high wtRAS activity in absence of high JAK2 phosphorylation levels might activate RAS via a different pathway yet to be uncovered for DS-ALL.

Taking RAS activity and inducibility, integrated with other protein activation patterns, we performed a multivariate analysis clustering that identified SR and HR groups for DS-ALL and showed that protein activation pattern is independently predictive of outcome using multivariate Cox regression.

Ultimately, patient-specific inhibitor combinations based on analyzed pathway activities should be part of future precision medicine approaches for HR-ALL groups. Ph-like ALL patients are already being studied for the combined effects of PI3K/mTOR and JAK/STAT inhibitor treatment(6).

The RAS inhibitor Salirasib used in our study disrupts the spatiotemporal localization of active RAS but requires relatively high concentrations, thus rendering it ineffective in Phase II clinical trials(39). Newer RAS-inhibitors like the RAS-mimetic Rigosertib, which blocks the RBD in RAS-effectors, as seen for bRAF and p110αPI3K in our PLA analysis on MUTZ-5 cells, could also allow blocking of both wt and mutant RAS and Rigosertib is currently being evaluated in a Phase III study for MDS/AML(40). Our data reveal that reducing RAS activity via inhibition of PTPN11 catalytical action may provide a functional alternative for ALL cells, while blocking the phosphorylation of PTPN11 via JAK inhibitors was not sufficient to prevent RAS activity, and concordantly with our mechanistic insight was also unable to block the direct interaction between PTPN11 and RAS. Our findings suggest that, depending on the patient’s protein activity profile, RAS inhibition should be considered in combination with PI3K/mTOR and/or JAK/STAT inhibitors to further augment clinical treatment. In particular in DS-ALL, this strategy might be applicable to most HR patients. However, based on our data, the focus should not lie on targeting mutant-RAS alone but also the inhibition of overstimulated RAS pathway activity in absence of RAS mutations.

Childhood leukemia in DS is distinguished by a relatively specific pattern of acquired mutation changes, for both AML(41-46) and ALL(20, 21, 47), and the reasons for this are not fully explained. More generally, people with DS have an unusual epidemiological pattern of malignancy: increased incidence and mortality for childhood leukemias of all types, but much decreased childhood and adult solid tumors(48, 49). Functional consequences of an increased dose of some chromosome 21 genes may play important roles(49). Increased propensity for early hematopoietic (both myeloid and lymphoid) cell fate can be influenced by trisomy of *RUNX1*(50, 51), whereas *ERG* trisomy is linked to skewing of cell fate towards megakaryocytic lineage (the most frequent AML form in DS)(52). Increased *HGMN1*-dose through trisomy 21 directly enhanced the early B-lymphocyte precursors, and could play a role as one of the initiating events in ALL leukemogenesis(53), whereas *CHAF1B* trisomy may increase the risk of AML(54). One of the most dose sensitive chromosome 21 genes known for multiple pathway de-regulations when copy number is increased, is *DYRK1A*. Increased *DYRK1A*-dose could promote both AML and ALL pathogenesis(55, 56). Interestingly, both *DYRK1A*, and another chromosome 21 gene *ITSN* also play a role in activating RAS, in specific cellular contexts(57, 58). It will be important to unravel the mechanisms behind the actions of these chromosome 21 genes, as their specific inhibition may be an additional component to consider in combinatorial therapy approaches(56). This is highlighted by very frequent observations of extra copies of chromosome 21 as acquired changes in DS and non-DS ALL, both at diagnosis, and at relapse(20, 59).

In conclusion, our data show that activation of RAS is a common feature of up to 80% of DS-ALL, and that patient pre-stratification for therapy optimization would be improved by assessing wtRAS protein activation status. Furthermore, our data suggest that inhibiting overstimulated RAS pathway activity should be a therapeutic strategy even in the absence of RAS mutations.

## Funding source

Singapore Ministry of Education Academic Research Funds Tier 2 grants (2015-T2-2-119 and 2015-T2-1-023); AIRC IG 19186 and Fondazione Cariparo 17/07 to G.B.

## Conflicts of Interest Disclosure

The authors declare no potential conflicts of interest.

## Author’s Contributions

D.K., D.R., and E.G. performed experiments; D.K., D.R., Z.L., E.G., J.G., I.A., S.K.K, W.J.C., H.A., D.M.W., A.E.Y., G.B., and D.N. analyzed results; D.K., Z.L., and D.N. prepared the figures; D.K., D.R., A.E.Y., G.B., and D.N. designed the research and wrote the paper.

## References

1. Inaba H, Greaves M, Mullighan CG. Acute lymphoblastic leukaemia. Lancet. 2013;381(9881):1943–55.

2. Pui CH, Robison LL, Look AT. Acute lymphoblastic leukaemia. Lancet. 2008;371(9617):1030–43.

3. Nguyen K, Devidas M, Cheng SC, La M, Raetz EA, Carroll WL, et al. Factors influencing survival after relapse from acute lymphoblastic leukemia: a Children’s Oncology Group study. Leukemia. 2008;22(12):2142–50.

4. Pui CH, Yang JJ, Hunger SP, Pieters R, Schrappe M, Biondi A, et al. Childhood Acute Lymphoblastic Leukemia: Progress Through Collaboration. J Clin Oncol. 2015;33(27):2938–48.

5. Bhojwani D, Pui CH. Relapsed childhood acute lymphoblastic leukaemia. Lancet Oncol. 2013;14(6):e205–17.

6. Tasian SK, Teachey DT, Li Y, Shen F, Harvey RC, Chen IM, et al. Potent efficacy of combined PI3K/mTOR and JAK or ABL inhibition in murine xenograft models of Ph-like acute lymphoblastic leukemia. Blood. 2017;129(2):177–87.

7. Hunger SP, Mullighan CG. Redefining ALL classification: toward detecting high-risk ALL and implementing precision medicine. Blood. 2015;125(26):3977.

8. Tasian SK, Loh ML, Hunger SP. Philadelphia chromosome–like acute lymphoblastic leukemia. Blood. 2017;130(19):2064–72.

9. Holmfeldt L, Wei L, Diaz-Flores E, Walsh M, Zhang J, Ding L, et al. The genomic landscape of hypodiploid acute lymphoblastic leukemia. Nat Genet. 2013;45(3):242–52.

10. Den Boer ML, van Slegtenhorst M, De Menezes RX, Cheok MH, Buijs-Gladdines JG, Peters ST, et al. A subtype of childhood acute lymphoblastic leukaemia with poor treatment outcome: a genome-wide classification study. Lancet Oncol. 2009;10(2):125–34.

11. Mullighan CG, Su X, Zhang J, Radtke I, Phillips LA, Miller CB, et al. Deletion of IKZF1 and prognosis in acute lymphoblastic leukemia. N Engl J Med. 2009;360(5):470–80.

12. Knight T, Irving JAE. Ras/Raf/MEK/ERK Pathway Activation in Childhood Acute Lymphoblastic Leukemia and Its Therapeutic Targeting. Frontiers in Oncology. 2014;4.

13. Ryan SL, Matheson E, Grossmann V, Sinclair P, Bashton M, Schwab C, et al. The role of the RAS pathway in iAMP21-ALL. Leukemia. 2016;30(9):1824–31.

14. Buitenkamp TD, Izraeli S, Zimmermann M, Forestier E, Heerema NA, van den Heuvel-Eibrink MM, et al. Acute lymphoblastic leukemia in children with Down syndrome: a retrospective analysis from the Ponte di Legno study group. Blood. 2014;123(1):70–7.

15. Lee P, Bhansali R, Izraeli S, Hijiya N, Crispino JD. The biology, pathogenesis and clinical aspects of acute lymphoblastic leukemia in children with Down syndrome. Leukemia. 2016;30(9):1816–23.

16. Harrison CJ. Targeting signaling pathways in acute lymphoblastic leukemia: new insights. Hematology Am Soc Hematol Educ Program. 2013;2013:118–25.

17. Russell LJ, Capasso M, Vater I, Akasaka T, Bernard OA, Calasanz MJ, et al. Deregulated expression of cytokine receptor gene, CRLF2, is involved in lymphoid transformation in B-cell precursor acute lymphoblastic leukemia. Blood. 2009;114(13):2688–98.

18. Roberts KG, Morin RD, Zhang J, Hirst M, Zhao Y, Su X, et al. Genetic alterations activating kinase and cytokine receptor signaling in high-risk acute lymphoblastic leukemia. Cancer Cell. 2012;22(2):153–66.

19. Roberts KG, Mullighan CG. Genomics in acute lymphoblastic leukaemia: insights and treatment implications. Nature Reviews Clinical Oncology. 2015;12(6):344–57.

20. Nikolaev SI, Garieri M, Santoni F, Falconnet E, Ribaux P, Guipponi M, et al. Frequent cases of RAS-mutated Down syndrome acute lymphoblastic leukaemia lack JAK2 mutations. Nature communications. 2014;5:4654.

21. Schwartzman O, Savino AM, Gombert M, Palmi C, Cario G, Schrappe M, et al. Suppressors and activators of JAK-STAT signaling at diagnosis and relapse of acute lymphoblastic leukemia in Down syndrome. Proceedings of the National Academy of Sciences of the United States of America. 2017;114(20):E4030–E9.

22. Yeoh AE, Ariffin H, Chai EL, Kwok CS, Chan YH, Ponnudurai K, et al. Minimal residual disease-guided treatment deintensification for children with acute lymphoblastic leukemia: results from the Malaysia-Singapore acute lymphoblastic leukemia 2003 study. J Clin Oncol. 2012;30(19):2384–92.

23. Gorban AN, Pitenko, A., Zinovyev, A. ViDaExpert: user-friendly tool for non-linear visualization and analysis of multidimensional vectorial data. 2014(28 May 2019).

24. Gaetano J. Holm-Bonferroni sequential correction: An EXCEL calculator (1.2) [Microsoft Excel workbook]. https://www.researchgatenet/publication/242331583_Holm-Bonferroni_Sequential_Correction_An_EXCEL_Calculator_-_Ver_12 2013.

25. Stangroom J. Fisher Exact Test calculator: Social Science Statistics; 2019 [updated 201928 May 2019]. Available from: https://www.socscistatistics.com/tests/fisher/default2.aspx.

26. Yoda A, Yoda Y, Chiaretti S, Bar-Natan M, Mani K, Rodig SJ, et al. Functional screening identifies CRLF2 in precursor B-cell acute lymphoblastic leukemia. Proceedings of the National Academy of Sciences of the United States of America. 2010;107(1):252–7.

27. Mohi MG, Arai K, Watanabe S. Activation and functional analysis of Janus kinase 2 in BA/F3 cells using the coumermycin/gyrase B system. Molecular biology of the cell. 1998;9(12):3299–308.

28. Barretina J, Caponigro G, Stransky N, Venkatesan K, Margolin AA, Kim S, et al. The Cancer Cell Line Encyclopedia enables predictive modelling of anticancer drug sensitivity. Nature. 2012;483(7391):603–7.

29. Tasian SK, Doral MY, Borowitz MJ, Wood BL, Chen IM, Harvey RC, et al. Aberrant STAT5 and PI3K/mTOR pathway signaling occurs in human CRLF2-rearranged B-precursor acute lymphoblastic leukemia. Blood. 2012;120(4):833–42.

30. Castellano E, Downward J. RAS Interaction with PI3K: More Than Just Another Effector Pathway. Genes Cancer. 2011;2(3):261–74.

31. Mendoza MC, Er EE, Blenis J. The Ras-ERK and PI3K-mTOR Pathways: Cross-talk and Compensation. Trends in biochemical sciences. 2011;36(6):320.

32. Hobbs GA, Der CJ, Rossman KL. RAS isoforms and mutations in cancer at a glance. J Cell Sci. 2016;129(7):1287–92.

33. McMahon LP, Yue W, Santen RJ, Lawrence JC, Jr. Farnesylthiosalicylic acid inhibits mammalian target of rapamycin (mTOR) activity both in cells and in vitro by promoting dissociation of the mTOR-raptor complex. Molecular endocrinology. 2005;19(1):175–83.

34. Bunda S, Burrell K, Heir P, Zeng L, Alamsahebpour A, Kano Y, et al. Inhibition of SHP2-mediated dephosphorylation of Ras suppresses oncogenesis. Nature communications. 2015;6:8859.

35. Schaper F, Gendo C, Eck M, Schmitz J, Grimm C, Anhuf D, et al. Activation of the protein tyrosine phosphatase SHP2 via the interleukin-6 signal transducing receptor protein gp130 requires tyrosine kinase Jak1 and limits acute-phase protein expression. The Biochemical journal. 1998;335 (Pt 3):557–65.

36. Roberts KG. Why and how to treat Ph-like ALL? Best Pract Res Clin Haematol. 2018;31(4):351–6.

37. Roberts KG, Yang YL, Payne-Turner D, Lin W, Files JK, Dickerson K, et al. Oncogenic role and therapeutic targeting of ABL-class and JAK-STAT activating kinase alterations in Ph-like ALL. Blood Adv. 2017;1(20):1657–71.

38. Potter N, Jones L, Blair H, Strehl S, Harrison CJ, Greaves M, et al. Single-cell analysis identifies CRLF2 rearrangements as both early and late events in Down syndrome and non-Down syndrome acute lymphoblastic leukaemia. Leukemia. 2018:1.

39. Riely GJ, Johnson ML, Medina C, Rizvi NA, Miller VA, Kris MG, et al. A phase II trial of Salirasib in patients with lung adenocarcinomas with KRAS mutations. J Thorac Oncol. 2011;6(8):1435–7.

40. Navada SC, Fruchtman SM, Odchimar-Reissig R, Demakos EP, Petrone ME, Zbyszewski PS, et al. A phase 1/2 study of rigosertib in patients with myelodysplastic syndromes (MDS) and MDS progressed to acute myeloid leukemia. Leukemia research. 2018;64:10–6.

41. De Vita S, Mulligan C, McElwaine S, Dagna-Bricarelli F, Spinelli M, Basso G, et al. Loss-of-function JAK3 mutations in TMD and AMKL of Down syndrome. Br J Haematol. 2007;137(4):337–41.

42. Groet J, McElwaine S, Spinelli M, Rinaldi A, Burtscher I, Mulligan C, et al. Acquired mutations in GATA1 in neonates with Down’s syndrome with transient myeloid disorder. Lancet. 2003;361(9369):1617–20.

43. Nikolaev SI, Santoni F, Vannier A, Falconnet E, Giarin E, Basso G, et al. Exome sequencing identifies putative drivers of progression of transient myeloproliferative disorder to AMKL in infants with Down syndrome. Blood. 2013;122(4):554–61.

44. Norton A, Fisher C, Liu H, Wen Q, Mundschau G, Fuster JL, et al. Analysis of JAK3, JAK2, and C-MPL mutations in transient myeloproliferative disorder and myeloid leukemia of Down syndrome blasts in children with Down syndrome. Blood. 2007;110(3):1077–9.

45. Vyas P, Roberts I. Down myeloid disorders: a paradigm for childhood preleukaemia and leukaemia and insights into normal megakaryopoiesis. Early Hum Dev. 2006;82(12):767–73.

46. Wechsler J, Greene M, McDevitt MA, Anastasi J, Karp JE, Le Beau MM, et al. Acquired mutations in GATA1 in the megakaryoblastic leukemia of Down syndrome. Nat Genet. 2002;32(1):148–52.

47. Bercovich D, Ganmore I, Scott LM, Wainreb G, Birger Y, Elimelech A, et al. Mutations of JAK2 in acute lymphoblastic leukaemias associated with Down’s syndrome. Lancet. 2008;372(9648):1484–92.

48. Hasle H, Friedman JM, Olsen JH, Rasmussen SA. Low risk of solid tumors in persons with Down syndrome. Genetics in medicine : official journal of the American College of Medical Genetics. 2016;18(11):1151–7.

49. Nizetic D, Groet J. Tumorigenesis in Down’s syndrome: big lessons from a small chromosome. Nature reviews Cancer. 2012;12(10):721–32.

50. De Vita S, Canzonetta C, Mulligan C, Delom F, Groet J, Baldo C, et al. Trisomic dose of several chromosome 21 genes perturbs haematopoietic stem and progenitor cell differentiation in Down’s syndrome. Oncogene. 2010;29(46):6102–14.

51. Lie ALM, Marinopoulou E, Lilly AJ, Challinor M, Patel R, Lancrin C, et al. Regulation of RUNX1 dosage is crucial for efficient blood formation from hemogenic endothelium. Development. 2018;145(5).

52. Salek-Ardakani S, Smooha G, de Boer J, Sebire NJ, Morrow M, Rainis L, et al. ERG is a megakaryocytic oncogene. Cancer Res. 2009;69(11):4665–73.

53. Lane AA, Chapuy B, Lin CY, Tivey T, Li H, Townsend EC, et al. Triplication of a 21q22 region contributes to B cell transformation through HMGN1 overexpression and loss of histone H3 Lys27 trimethylation. Nat Genet. 2014;46(6):618–23.

54. Volk A, Liang K, Suraneni P, Li X, Zhao J, Bulic M, et al. A CHAF1B-Dependent Molecular Switch in Hematopoiesis and Leukemia Pathogenesis. Cancer Cell. 2018;34(5):707–23 e7.

55. Malinge S, Bliss-Moreau M, Kirsammer G, Diebold L, Chlon T, Gurbuxani S, et al. Increased dosage of the chromosome 21 ortholog Dyrk1a promotes megakaryoblastic leukemia in a murine model of Down syndrome. J Clin Invest. 2012;122(3):948–62.

56. Thompson BJ, Bhansali R, Diebold L, Cook DE, Stolzenburg L, Casagrande AS, et al. DYRK1A controls the transition from proliferation to quiescence during lymphoid development by destabilizing Cyclin D3. J Exp Med. 2015;212(6):953–70.

57. Kelly PA, Rahmani Z. DYRK1A enhances the mitogen-activated protein kinase cascade in PC12 cells by forming a complex with Ras, B-Raf, and MEK1. Molecular biology of the cell. 2005;16(8):3562–73.

58. Mohney RP, Das M, Bivona TG, Hanes R, Adams AG, Philips MR, et al. Intersectin activates Ras but stimulates transcription through an independent pathway involving JNK. J Biol Chem. 2003;278(47):47038–45.

59. Berger R. Acute lymphoblastic leukemia and chromosome 21. Cancer Genet Cytogenet. 1997;94(1):8–12.

