## Supplementary material for "RAS activation via CRLF2 signaling is a widespread mechanism in Down syndrome acute lymphoblastic leukemia regardless of RAS mutations"

**Supplemental material**

**Supplementary methods**

**Ba/F3 cells transduction**

Ba/F3 cells were nucleofected using a Nucleofector 2b (Lonza, Basel, CH) using Cell Line Nucleofector Kit V (Lonza) according to the manufacturer's protocol. The plasmids used for Ba/F3 transfection were MSCVpuromycinR-hCRLF2 (human wt CRLF2) and MSCVneomycinR-hJAK2RG (human $JAK2_{R683G}$ mutant)(1). In order to select for stably transfected cells, MSCVpuromycinR-hCRLF2-transfected cells were cultured with 1 µg/mL puromycin (Cat.#P8833; Sigma-Aldrich), MSCVneomycinR-hJAK2RG-transfected cells were cultured with 1.8 mg/mL G418 (Cat.#108321-42-2; Sigma-Aldrich), and co-transfected cells were cultured with both antibiotics together. Cells transfected with the plasmid pMax-EGFP (Lonza) were used as control.

**DNA- and RNA-sequencing of primary patient material**

Patient samples were either whole exome-sequenced previously(2) or amplicons of *JAK2*, *KRAS* and *NRAS* (in addition, *HRAS* of MUTZ-5 cells) were amplified from genomic DNA and sent for Sanger sequencing (1st BASE, Singapore, SG) in both orientations and screened for mutations in Mutation Surveyor v5.0.1 (Softgenetics, State College, US). Oligonucleotides (Supplementary-Tab.S3) were designed to cover known activating mutation hotspots.

RNA sequencing was performed for patients of the MS2003/2010 studies(3, 4) using TruSeq Stranded mRNA Library Prep kit (Illumina, San Diego, US); sequenced on HiSeq2000/2500 or NextSeq500 (Illumina). Reads were aligned to hg19-reference genome using Tophat2(5). The number of reads mapped to each gene were counted using featureCounts(6) and gene expression level was calculated as fragments per kilobase of transcript per million mapped reads (FPKM). GATK best practices was performed for variant calling on RNAseq(7). Data was submitted to the European Genome-phenome Archive (Accession-number EGAS00001001858).

**Immunofluorescence**

MUTZ-5 cells were fixated with 4% paraformaldehyde (PFA; Cat.#30525-89-4/158127; Sigma-Aldrich), permeabilized with methanol (Cat.#67-56-1; Sigma-Aldrich) at -20 °C and then stained with primary antibodies against either pan-RAS or bRAF. Secondary antibodies, labeled with either Alexa-488 (green) or Alexa-594 (red), specific to the respective primary antibody species were used to visualize the spatial organization of RAS (mostly at plasma membrane) and RAF (mostly in cytoplasma) proteins in a LSM800 inverted confocal microscope with Airyscan, 63×/NA1.20W objective (Carl Zeiss, Oberkochen, DE) at room temperature. Cell nuclei were stained with DAPI. Images were captured using ZEN software (Carl Zeiss) and processed in ImageJ.

**Quantitative PCR**

Quantitative PCR was performed using a StepOnePlus Real-Time PCR system (Applied Biosystems, Foster City, USA) with Power SYBR Green PCR master mix (Applied Biosystems) according to the standard manufacturer’s protocol. The expression of human CRLF2 or human JAK2 was standardized against the expression of mouse GAPDH in each sample. The qPCR oligonucleotides are listed in Supplementary-Tab.S3.

**Supplementary results**

**(Expanded results description) Higher levels of RAS protein and mRNA correlate with poor outcome in DS-ALL and non-DS childhood ALL**

The entire analysis shown so far was based on differences in activity of proteins targets. We also wanted to address protein expression levels in DS-ALL SR and HR groups. Based on individual protein expression levels (Supplementary-Fig.S6A), we found that overall levels of both RAS and rpS6 positively correlated with HR, but levels of CRLF2, STAT5, JAK2, MEK1/2, or ERK1/2 did not. In order to examine similar correlations in non-DS ALL, the material availability only permitted us to look for differences in mRNA transcript levels of the same genes within the whole transcriptome RNA-seq data of the MS2003/2010 cohort (N=346 non-DS ALL cases; median follow-up via reverse Kaplan-Meier = 7.64 years)(3, 4). *CRLF2* mRNA expression (Supplementary-Fig.S6B) and Cox regression hazard ratio (Supplementary-Fig.S7A) were significantly increased in the poor first event samples (n=47). We focused all further analysis on non-DS B-ALL samples with high-risk genetics: Moderate or high *CRLF2* expression (log2(FPKM+1)>0.7) and excluded ALL-subtypes that are considered to confer favorable outcomes (ETV6-RUNX1, Hyperdiploid, TCF3-PBX1, DUX4 and ZNF384). Within the resulting HR sub-cohort (n=91; median follow-up = 6.95 years) KRAS mRNA-expression was independently predictive of outcome (*P*=.030) when included in a multivariate analysis using Cox proportional hazards regression model together with Ph-like status, RAS mutation status, sex, and NCI risk (Supplementary-Fig.S7B). Of all analyzed genes, exclusively KRAS and JAK2 mRNA expression correlated with poor outcome (Supplementary-Fig.S6B). However, only high KRAS mRNA-levels were also significant in Kaplan–Meier survival estimator (Supplementary-Fig.S7B).

This provides just a hint of a trend compatible with conclusions reached for the general role of RAS (irrespective of its mutational status) as a biomarker in childhood ALL, but emphasizes the need to examine the protein-activation and inducibility parameters on a much larger cohort of non-DS ALL patients for a more accurate risk-prediction sub-stratification.

**Supplementary discussion**

**(Expanded) RAS inhibition strategies**

The importance of identifying RAS activity for the patient outcome relies on the availability of effective RAS-treatments. RAS has been commonly deemed to be ‘undruggable’, a term describing the lack of RAS inhibitors that perform well pharmacologically. The RAS inhibitor Salirasib, a Farnesyl Thiosalicylic Acid (FTS) acts non-direct, as a mimetic of RAS for binding to RAS-escort proteins which selectively disrupts the association of RAS to the plasma membrane. It showed no effect in clinical trials on solid cancers trials(8) and requires relatively high dosage. We chose this RAS inhibitor for being able to use a single inhibitor that can block RAS activity of all the main RAS isoforms and most importantly can also block wtRas activity independent of RAS mutations. A new generation of RAS inhibitors are on the horizon(9) such as mutant-specific inhibitors of KRAS(G12C)(10). However, based on our data, the focus should not lie on targeting mutant-RAS alone but also the inhibition of overstimulated activity in absence of RAS mutations.

The RAS-mimetic Rigosertib allows to block the interaction between RAS family members and their numerous effector proteins by hijacking the RBD in RAS-effector proteins and showed promising results in a Phase II study and is currently being evaluated in a Phase III study for MDS/AML(11). Treatments involving the inhibition of activated RAS, independent of mutation status, that block the activation of multiple RAS-effector pathways, could help to cripple the cancer cells’ ability to adapt.

**Supplementary references**

1. van Bodegom D, Zhong J, Kopp N, Dutta C, Kim MS, Bird L, et al. Differences in signaling through the B-cell leukemia oncoprotein CRLF2 in response to TSLP and through mutant JAK2. Blood. 2012;120(14):2853-63.
2. Nikolaev SI, Garieri M, Santoni F, Falconnet E, Ribaux P, Guipponi M, et al. Frequent cases of RAS-mutated Down syndrome acute lymphoblastic leukaemia lack JAK2 mutations. Nature communications. 2014;5:4654.
3. Yeoh AE, Ariffin H, Chai EL, Kwok CS, Chan YH, Ponnudurai K, et al. Minimal residual disease-guided treatment deintensification for children with acute lymphoblastic leukemia: results from the Malaysia-Singapore acute lymphoblastic leukemia 2003 study. J Clin Oncol. 2012;30(19):2384-92.
4. Yeoh AEJ, Lu Y, Chin WHN, Chiew EKH, Lim EH, Li Z, et al. Intensifying Treatment of Childhood B-Lymphoblastic Leukemia With IKZF1 Deletion Reduces Relapse and Improves Overall Survival: Results of Malaysia-Singapore ALL 2010 Study. J Clin Oncol. 2018;36(26):2726-35.
5. Kim D, Pertea G, Trapnell C, Pimentel H, Kelley R, Salzberg SL. TopHat2: accurate alignment of transcriptomes in the presence of insertions, deletions and gene fusions. Genome Biol. 2013;14(4):R36.
6. Liao Y, Smyth GK, Shi W. featureCounts: an efficient general purpose program for assigning sequence reads to genomic features. Bioinformatics. 2014;30(7):923-30.
7. Van der Auwera GA, Carneiro MO, Hartl C, Poplin R, Del Angel G, Levy-Moonshine A, et al. From FastQ data to high confidence variant calls: the Genome Analysis Toolkit best practices pipeline. Curr Protoc Bioinformatics. 2013;43:11 0 1-33.
8. Riely GJ, Johnson ML, Medina C, Rizvi NA, Miller VA, Kris MG, et al. A phase II trial of Salirasib in patients with lung adenocarcinomas with KRAS mutations. Journal of Thoracic Oncology: Official Publication of the International Association for the Study of Lung Cancer. 2011;6(8):1435-7.
9. Dang CV, Reddy EP, Shokat KM, Soucek L. Drugging the 'undruggable' cancer targets. Nature reviews Cancer. 2017;17(8):502-8.
10. Patricelli MP, Janes MR, Li LS, Hansen R, Peters U, Kessler LV, et al. Selective Inhibition of Oncogenic KRAS Output with Small Molecules Targeting the Inactive State. Cancer discovery. 2016;6(3):316-29.
11. Navada SC, Fruchtman SM, Odchimar-Reissig R, Demakos EP, Petrone ME, Zbyszewski PS, et al. A phase 1/2 study of rigosertib in patients with myelodysplastic syndromes (MDS) and MDS progressed to acute myeloid leukemia. Leukemia research. 2018;64:10-6.
12. Yoda A, Yoda Y, Chiaretti S, Bar-Natan M, Mani K, Rodig SJ, et al. Functional screening identifies CRLF2 in precursor B-cell acute lymphoblastic leukemia. Proceedings of the National Academy of Sciences of the United States of America. 2010;107(1):252-7.

**Supplementary-Tab.S1) Clinical and biological characteristics of the leukemia samples**

| DS-ALL Sample ID | Presentation / Relapse | BM/PB | AIEOP-BFM subclassification | Blasts % | WBC | Diagnosis | Karyotype | Gender | NCI | Poor outcome (death or relapse) | Sample ID in: https://doi-org.ezlibproxy1.ntu.edu.sg/10.1038/ncomms5654 | |
|---|---|---|---|---|---|---|---|---|---|---|---|---|
| DS01 | Presentation | BM | B-II | 65.0 | N/A | ALL | N.D. | M | N/A | N | 4-23-T1 | |
| DS02 | Presentation | BM | B-II | 90.0 | 122800 | ALL | N.D. | M | HR | Y | | |
| DS04 | Presentation | BM | B-II | 95.0 | N/A | ALL | 47,XY,+21c | M | SR | N | 4-1030604-T1 | |
| DS05 | Presentation | BM | B-II | 95.0 | N/A | ALL | N.D. | M | HR | N | | |
| DS06 | Presentation | BM | B-III | 91.6 | 323000 | ALL | N.D. | F | HR | N | 4-37-T1 | |
| DS07 | Presentation | BM | B-II | 87.0 | 86400 | ALL | N.D. | F | HR | Y | | |
| DS08 | Presentation | BM | B-II | 85.0 | N/A | ALL | N.D. | F | SR | N | | |
| DS09 | Presentation | BM | B-II | 70.0 | 18780 | ALL | 47,XY,+21c | M | SR | Y | 4-1036101-T1 | |
| DS10 | Presentation | BM | B-II | 62.0 | 35400 | ALL | 47,XY,+21c | M | HR | Y | 4-1036272-T1 | |
| DS11 | Presentation | BM | B-III | 91.0 | N/A | ALL | 48,XY,+X,+21 | M | SR | Y | | |
| DS16 | Presentation | BM | B-II | 90.0 | 2400 | ALL | N.D. | M | HR | Y | 4-44-T1 | |
| DS17 | Presentation | BM | B-II | 90.0 | 44600 | ALL | 47,XY,+21c[1 | M | HR | Y | 4-03-T1 | |
| DS18 | Presentation | BM | B-II | 86.0 | 23530 | ALL | N.D. | M | SR | N | 4-02-T1 | |
| DS20 | Presentation | BM | B-II | 80.0 | 11500 | ALL | 47,XX,t(8;14 | F | SR | N | 4-29-T1 | also Remission sample tested |
| DS22 | Presentation | BM | B-II | 97.0 | N/A | ALL | N.D. | F | NA | Y | | |
| DS23 | Presentation | BM | B-II | 88.0 | 27400 | ALL | 47,XX,+21c | F | SR | N | 4-1044929-T1 | |
| DS26 | Presentation | BM | B-II | 88.0 | N/A | ALL | 47,XX,+21c[1 | F | N/A | N | | |
| DS27 | Presentation | N/A | N/A | N/A | N/A | ALL | N/A | M | N/A | N | | also Remission sample tested |
| DS29 | Presentation | BM | B-II | 87.0 | 55000 | ALL | 47,XY,+21c[1 | M | HR | Y | | |
| DS30 | Presentation | PB | B-II | 82.0 | 206000 | ALL | N.D. | F | HR | N | | |
| DS09R | Relapse | BM | B-II | 47.0 | 12200 | ALL | 47,XY,+21c | M | | | 4-1036101-T2 | |
| DS12R | Relapse | BM | B-II | 79.0 | 17000 | ALL | N.D. | M | | | | |
| DS16R | Relapse | BM | B-II | 90.0 | N/A | ALL | N.D. | M | | | 4-44-T2 | |
| DS19R | Relapse | BM | B-II | 88.0 | 221400 | ALL | N.D. | M | | | | |
| DS22R | Relapse | BM | B-III | 87.0 | 46000 | ALL | N.D. | F | | | 4-29-T2 | |
| DS28R | Relapse | BM | B-II | 83.0 | 98400 | ALL | N.D. | M | | | | |
| DS29R | Relapse | BM | B-II | 83.0 | 39800 | ALL | N.D. | M | | | | |
| DS24m | Remission | BM | N/A | N/A | N/A | N/A | N.D. | M | | Y | | |
| DS25m | Remission | BM | N/A | N/A | N/A | N/A | N.D. | F | | Y | | |

| Non-DS ALL Sample ID | Presentation / Relapse | BM/PB | AIEOP-BFM subclassification | Blasts % | WBC | Diagnosis | Karyotype | Gender | NCI | Poor outcome (death or relapse) |
|---|---|---|---|---|---|---|---|---|---|---|
| NDS03 | Presentation | N/A | N/A | N/A | N/A | ALL | N/A | F | N/A | N |
| NDS04 | Presentation | BM | B-II | 93 | 97800 | ALL | N.D. | M | HR | N |
| NDS05 | Presentation | BM | B-II | 82.8 | 56000 | ALL | 46,XX | F | HR | Y |
| NDS06 | Presentation | BM | B-II | 89 | 80000 | ALL | 46,XX | F | HR | N |

| | | | | | | | | |
|---|---|---|---|---|---|---|---|---|
| NDS01R | Relapse | BM | B-II | 44.1 | NA | ALL | N.D. | F |
| NDS02R | Relapse | BM | B-II | 97 | NA | ALL | N.D. | M |

**Supplementary-Tab.S1) Clinical and biological characteristics of the leukemia samples used in this study and therapy outcomes.**

**Supplementary-Tab.S2) List of Bonferroni-corrected *P* -values**

**Fig.1B)**

| Comparison: | Bonferroni p-value after one-way ANOVA: |
|---|---|
| — vs. hJAK2$^{R683G}$ | >0.99999 |
| — vs. hCRLF2 | >0.99999 |
| — vs. hJAK2$^{R683G}$ + hCRLF2 | 0.00160 |
| hJAK2$^{R683G}$ vs. hCRLF2 | >0.99999 |
| hJAK2$^{R683G}$ vs. hJAK2$^{R683G}$ + hCRLF2 | 0.00364 |
| hCRLF2 vs. hJAK2$^{R683G}$ + hCRLF2 | 0.00347 |

**Fig.2A)**

| Comparison: | p-value: | Holm-Bonferroni corrected p value: |
|---|---|---|
| JAK2 activity uninduced vs. TSLP | 0.00043 | 0.00171 |
| MEK1/2 activity uninduced vs. TSLP | 0.00065 | 0.00195 |
| Pan-Ras activity uninduced vs. TSLP | 0.00308 | 0.00616 |
| PTPN11 activity uninduced vs. TSLP | 0.01846 | 0.01846 |

**Fig.2C)**

| Comparison: | p-value: | Holm-Bonferroni corrected p value: |
|---|---|---|
| KRas activity uninduced vs. TSLP | 0.00216 | 0.00648 |
| NRas activity uninduced vs. TSLP | 0.01905 | 0.03810 |
| HRas activity uninduced vs. TSLP | 0.06632 | 0.06632 |

**Fig.4B)**

| Comparison: | Bonferroni p-value after one-way ANOVA: |
|---|---|
| Uninduced (+DMSO) vs. TSLP (+DMSO) | 0.01572 |
| Uninduced (+DMSO) vs. TSLP + Ras inh. | >0.99999 |
| Uninduced (+DMSO) vs. TSLP + JAK inh. | 0.20936 |
| Uninduced (+DMSO) vs. PLA negative control | 0.04445 |
| TSLP (+DMSO) vs. TSLP + Ras inh. | 0.02177 |
| TSLP (+DMSO) vs. TSLP + JAK inh. | 0.00214 |
| TSLP (+DMSO) vs. PLA negative control | 0.00102 |
| TSLP + Ras inh. vs. TSLP + JAK inh. | 0.13163 |
| TSLP + Ras inh. vs. PLA negative control | 0.03079 |
| TSLP + JAK inh. vs. PLA negative control | >0.99999 |

**Fig.5A)**

| Comparison: | Bonferroni p-value after one-way ANOVA: |
|---|---|
| DMSO uninduced vs. DMSO TSLP | >0.99999 |
| DMSO uninduced vs. Ras inh. uninduced | <0.00001 |
| DMSO uninduced vs. Ras inh. TSLP | <0.00001 |
| DMSO uninduced vs. PI3K inh. uninduced | <0.00001 |
| DMSO uninduced vs. PI3K inh. TSLP | <0.00001 |
| DMSO uninduced vs. JAK inh. uninduced | >0.99999 |
| DMSO uninduced vs. JAK inh. TSLP | 0.33322 |
| DMSO TSLP vs. Ras inh. uninduced | <0.00001 |
| DMSO TSLP vs. Ras inh. TSLP | <0.00001 |
| DMSO TSLP vs. PI3K inh. uninduced | <0.00001 |
| DMSO TSLP vs. PI3K inh. TSLP | <0.00001 |
| DMSO TSLP vs. JAK inh. uninduced | >0.99999 |
| DMSO TSLP vs. JAK inh. TSLP | 0.00589 |
| Ras inh. uninduced vs. Ras inh. TSLP | >0.99999 |
| Ras inh. uninduced vs. PI3K inh. uninduced | 0.00111 |
| Ras inh. uninduced vs. PI3K inh. TSLP | >0.99999 |
| Ras inh. uninduced vs. JAK inh. uninduced | <0.00001 |
| Ras inh. uninduced vs. JAK inh. TSLP | 0.00006 |
| Ras inh. TSLP vs. PI3K inh. uninduced | <0.00001 |
| Ras inh. TSLP vs. PI3K inh. TSLP | 0.31820 |
| Ras inh. TSLP vs. JAK inh. uninduced | <0.00001 |
| Ras inh. TSLP vs. JAK inh. TSLP | 0.02665 |
| PI3K inh. uninduced vs. PI3K inh. TSLP | 0.00944 |
| PI3K inh. uninduced vs. JAK inh. uninduced | <0.00001 |

**Fig.3A**

| Comparison: | p-value: | Holm-Bonferroni corrected p value: |
|---|---|---|
| RAS : p-PTPN11 | 0.001306 | 0.002612 |
| NC (rpS6 : p-bRAF | 0.956956 | >0.999999 |

**Fig.3C**

| Comparison: | Bonferroni p-value after one-way ANOVA: |
|---|---|
| DMSO vs. DMSO + | 0.00748 |
| DMSO vs. PTPN11 | 0.01592 |
| DMSO + TSLP vs. | 0.00027 |

**Fig.7D)**

| Comparison: | p-value: | Holm-Bonferroni corrected p value: |
|---|---|---|
| Pan-Ras basal activity SR vs HR | 0.01315 | 0.03165 |
| JAK2 basal activity SR vs HR | 0.00053 | 0.00265 |
| STAT5 basal activity SR vs HR | 0.55617 | 0.55617 |
| MEK basal activity SR vs HR | 0.00495 | 0.01980 |
| ERK basal activity SR vs HR | 0.00035 | 0.00209 |
| S6 basal activity SR vs HR | 0.01055 | 0.03165 |
| Pan-Ras TSLP-induced activity SR vs HR | 0.00383 | 0.00766 |
| JAK2 TSLP-induced activity SR vs HR | 0.00027 | 0.00161 |
| STAT5 TSLP-induced activity SR vs HR | 0.00241 | 0.00722 |
| MEK TSLP-induced activity SR vs HR | 0.01622 | 0.01622 |
| ERK TSLP-induced activity SR vs HR | 0.00163 | 0.00653 |
| S6 TSLP-induced activity SR vs HR | 0.00032 | 0.00162 |

**Fig.8B)**

| Comparison: | Bonferroni p-value after one-way ANOVA: |
|---|---|
| **Pan-Ras activity** | |
| pan-Ras inh. vs. PI3K/mTOR dual inh. (PI-103) | 0.203 |
| pan-Ras inh. vs. JAK inh. (Ruxolitinib) | 0.203 |
| PI3K/mTOR dual inh. (PI-103) vs. JAK inh. (Ruxolitinib) | 0.893 |

| | |
|---|---|
| **JAK2 activity** | |
| pan-Ras inh. vs. PI3K/mTOR dual inh. (PI-103) | 0.910 |
| pan-Ras inh. vs. JAK inh. (Ruxolitinib) | 0.910 |
| PI3K/mTOR dual inh. (PI-103) vs. JAK inh. (Ruxolitinib) | 0.910 |

| | |
|---|---|
| **STAT5 activity** | |
| pan-Ras inh. vs. PI3K/mTOR dual inh. (PI-103) | 0.650 |
| pan-Ras inh. vs. JAK inh. (Ruxolitinib) | <.001 |
| PI3K/mTOR dual inh. (PI-103) vs. JAK inh. (Ruxolitinib) | <.001 |

| | |
|---|---|
| **MEK1/2 activity** | |
| pan-Ras inh. vs. PI3K/mTOR dual inh. (PI-103) | 0.572 |
| pan-Ras inh. vs. JAK inh. (Ruxolitinib) | 0.223 |
| PI3K/mTOR dual inh. (PI-103) vs. JAK inh. (Ruxolitinib) | 0.572 |

| | |
|---|---|
| **ERK1/2 activity** | |
| pan-Ras inh. vs. PI3K/mTOR dual inh. (PI-103) | 0.611 |
| pan-Ras inh. vs. JAK inh. (Ruxolitinib) | 0.266 |
| PI3K/mTOR dual inh. (PI-103) vs. JAK inh. (Ruxolitinib) | 0.611 |

**Supplementary-Fig.S1C)**

| Comparison: | Bonferroni p-value after one-way ANOVA: |
|---|---|
| **JAK2 mRNA levels:** | |

**Supplementary-Fig.S2C,D,E)**

| Comparison: | p-value: | Holm-Bonferroni corrected p value: |
|---|---|---|
| GRB2 : phospho-PTPN11 (uninducedvsTSLP) | 4.0599E-06 | 0.000434405 |

**Supplementary-Fig.S6A)**

| Comparison: | p-value: | Holm-Bonferroni corrected p value: |
|---|---|---|
| Pan-Ras protein expression SR vs. HR | 0.00427 | 0.02564 |

| | |
|---|---|
| — vs. hJAK2$^{R683G}$ | 0.00415 |
| — vs. hCRLF2 | >0.99999 |
| — vs. hJAK2$^{R683G}$ + hCRLF2 | 0.01031 |
| hJAK2$^{R683G}$ vs. hCRLF2 | 0.00413 |
| hJAK2$^{R683G}$ vs. hJAK2$^{R683G}$ + hCRLF2 | 0.69131 |
| hCRLF2 vs. hJAK2$^{R683G}$ + hCRLF2 | 0.01023 |

**Supplementary-Fig.S1D)**

| **CRLF2 mRNA levels:** | |
|---|---|
| — vs. hJAK2$^{R683G}$ | >0.99999 |
| — vs. hCRLF2 | 0.02509 |
| — vs. hJAK2$^{R683G}$ + hCRLF2 | 0.00020 |
| hJAK2$^{R683G}$ vs. hCRLF2 | 0.02514 |
| hJAK2$^{R683G}$ vs. hJAK2$^{R683G}$ + hCRLF2 | 0.00020 |
| hCRLF2 vs. hJAK2$^{R683G}$ + hCRLF2 | 0.00077 |

**Supplementary-Fig.S1E)**

| **Comparison:** | **Bonferroni p-value after one-way ANOVA:** |
|---|---|
| **Day4:** | |
| — vs. hJAK2$^{R683G}$ | >0.99999 |

| | | |
|---|---|---|
| pan-RAS : bRAF (TSLPvsRigo) | 8.1576E-05 | 0.006362897 |
| pan-RAS : p110aPI3K (TSLPvsRigo) | 0.0001618 | 0.0118114 |
| pan-RAS : phospho-PTPN11 (TSLPvsRigo) | 0.10487214 | 0.97912792 |
| pan-RAS : phospho-PTPN11 (unindurcedvsTSLP ) | 1.6756E-07 | 2.19501E-05 |
| pan-RAS : PTPN11 (TSLPvsRuxo) | 7.0237E-06 | 0.000709391 |
| pan-RAS : SOS1 (unindurcedvsTSLP ) | 0.0002915 | 0.019530583 |
| rpS6 : phospho-bRAF (TSLPvsRigo) | 3.5584E-06 | 0.000387867 |
| rpS6 : phospho-bRAF (TSLPvsRuxo) | 0.00040452 | 0.025484468 |
| rpS6 : phospho-bRAF (unindurcedvsTSLP ) | 0.00147548 | 0.070801112 |
| rpS6 : phospho-rpS6 (TSLPvsRigo) | 2.0311E-05 | 0.001828029 |
| rpS6 : phospho-rpS6 (TSLPvsRuxo) | 9.8818E-08 | 1.30439E-05 |
| rpS6 : phospho-rpS6 (unindurcedvsTSLP ) | 3.3757E-07 | 4.28713E-05 |
| SOS1 : GRB2 (unindurcedvsTSLP ) | 0.00023351 | 0.016345942 |

**Supplementary-Fig.S3B)**

| **Comparison:** | **p-value:** | **Holm-Bonferroni corrected p value:** |
|---|---|---|
| STAT5 activity uninduced vs. TSLP | 0.00006 | 0.00039 |
| JAK2 activity uninduced vs. TSLP | 0.00043 | 0.00256 |

| | |
|---|---|
| PI3K inh. uninduced vs. JAK inh. TSLP | <0.00001 |
| PI3K inh. TSLP vs. JAK inh. uninduced | <0.00001 |
| PI3K inh. TSLP vs. JAK inh. TSLP | <0.00001 |
| JAK inh. uninduced vs. JAK inh. TSLP | 0.11829 |

| | | |
|---|---|---|
| JAK2 protein expression SR vs. HR | 0.07137 | 0.28547 |
| STAT5 protein expression SR vs. HR | 0.14639 | 0.43916 |
| MEK1/2 protein expression SR vs. HR | 0.40022 | 0.80045 |
| ERK1/2 protein expression SR vs. HR | 0.04039 | 0.20196 |
| rpS6 protein expression SR vs. HR | 0.00034 | 0.00237 |
| CRLF2 protein expression SR vs. HR | 0.49585 | 0.80045 |

**Supplementary-Fig.S6B)**

| **Comparison:** | **p-value:** | **Holm-Bonferroni corrected p value:** |
|---|---|---|
| KRAS mRNA good vs. poor outcome | 0.02670 | 0.18693 |
| JAK2 mRNA good vs. poor outcome | 0.03945 | 0.23670 |
| NRAS/KRAS mRNA good vs. poor outcome | 0.04616 | 0.23670 |
| STAT5B mRNA good vs. poor outcome | 0.10881 | 0.43525 |
| RPS6 mRNA good vs. poor outcome | 0.81850 | >0.99999 |
| MEK1/2 mRNA good vs. poor outcome | 0.87458 | >0.99999 |
| ERK1/2 mRNA good vs. poor outcome | 0.95096 | >0.99999 |

| **rpS6 activity** | |
|---|---|
| pan-Ras inh. vs. PI3K/mTOR dual inh. (PI-103) | 0.237 |
| pan-Ras inh. vs. JAK inh. (Ruxolitinib) | 0.237 |
| PI3K/mTOR dual inh. (PI-103) vs. JAK inh. (Ruxolitinib) | 0.885 |

| **Measurement:** | **Comparison:** | **p-value:** | **Holm-Bonferroni corrected p value:** |
|---|---|---|---|
| STAT5 phosph. | JAK inh. vs. vehicle control | 0.00000 | 0.00001 |
| Pan-Ras-GTP level | Ras inh. vs. vehicle control | 0.00004 | 0.00061 |
| rpS6 phosph. | PI3K inh. vs. vehicle control | 0.00062 | 0.00988 |
| ERK1/2 phosph. | JAK inh. vs. vehicle control | 0.00213 | 0.03193 |
| rpS6 phosph. | JAK inh. vs. vehicle control | 0.00231 | 0.03231 |
| rpS6 phosph. | Ras inh. vs. vehicle control | 0.00320 | 0.04154 |
| MEK1/2 phosph. | JAK inh. vs. vehicle control | 0.00500 | 0.06004 |
| MEK1/2 phosph. | PI3K inh. vs. vehicle control | 0.00737 | 0.08112 |
| Pan-Ras-GTP level | JAK inh. vs. vehicle control | 0.01001 | 0.10007 |
| JAK2 phosph. | Ras inh. vs. vehicle control | 0.03552 | 0.31971 |
| JAK2 phosph. | PI3K inh. vs. vehicle control | 0.05956 | 0.47652 |
| JAK2 phosph. | JAK inh. vs. vehicle control | 0.06092 | 0.47652 |
| ERK1/2 phosph. | PI3K inh. vs. vehicle control | 0.06835 | 0.47652 |
| Pan-Ras-GTP level | PI3K inh. vs. vehicle control | 0.28436 | >0.99999 |
| MEK1/2 phosph. | Ras inh. vs. vehicle control | 0.46249 | >0.99999 |
| STAT5 phosph. | Ras inh. vs. vehicle control | 0.56159 | >0.99999 |

| | |
|---|---|
| — vs. hCRLF2 | 0.98323 |
| — vs. hJAK2$^{R683G}$ + hCRLF2 | 0.00016 |
| hJAK2$^{R683G}$ vs. hCRLF2 | 0.98323 |
| hJAK2$^{R683G}$ vs. hJAK2$^{R683G}$ + hCRLF2 | 0.00016 |
| hCRLF2 vs. hJAK2$^{R683G}$ + hCRLF2 | 0.00022 |

| | | |
|---|---|---|
| MEK activity uninduced vs. TSLP | 0.00065 | 0.00325 |
| rpS6 activity uninduced vs. TSLP | 0.00067 | 0.00325 |
| Pan-Ras activity uninduced vs. TSLP | 0.00308 | 0.01053 |
| PTPN11 activity uninduced vs. TSLP | 0.01846 | 0.03692 |
| ERK activity uninduced vs. TSLP | 0.02192 | 0.03692 |

| | | | |
|---|---|---|---|
| ERK1/2 phosph. | Ras inh. vs. vehicle control | 0.63041 | >0.99999 |
| STAT5 phosph. | PI3K inh. vs. vehicle control | 0.84173 | >0.99999 |

**Supplementary-Tab.S2) List of Bonferroni-adjusted or Holm-Bonferroni-corrected *P*-values from all statistical tests engaging in multiple comparisons in this article.** Original *P*-values before Holm-Bonferroni-correction are also listed; the P-values appearing in figures and supplementary figures are highlighted red. See Materials and Methods for more details about the statistical methods used.

**Supplementary-Tab.S3) List of oligonucleotides**

**human JAK2 sequencing:**

| Name: | Sequence: |
|---|---|
| Fwd-JAK2amp1,Seq1 | CCACTCTTGCTCTCTCTCACTTT |
| Fwd-JAK2amp2,Seq2 | CCATCCAGAAACACAAACCATGT |
| Fwd-JAK2amp3,Seq3 | CGCCTGTATTCCCAGCTACT |
| Fwd-JAK2,Seq4 | GGAAGTATTTGGCTAGTTGGT |
| Rev-JAK2amp3,rSeq1 | CCTAATGTCTACTTCAACACGGT |
| Rev-JAK2,rSeq2 | GGCAGCTTACCAGCACTGTA |
| Rev-JAK2amp2,rSeq3 | CCTTTACACCACTGCCCAAGTAA |
| Rev-JAK2amp1,rSeq4 | CCTAGCTGTGATCCTGAAACTGA |

**human KRAS sequencing:**

| Name: | Sequence: |
|---|---|
| Fwd KRas amplicon 1 | GCGTCGATGGAGGAGTTTGTA |
| Rev KRas amplicon 1 | GCACAGAGAGTGAACATCATGGA |
| Fwd KRas amplicon 2 | CCACCAGCAATGCACAAAGATT |
| Rev KRas amplicon 2 | CCAAAGCCAAAAGCAGTACCATG |
| Fwd KRas Seq3 | GGTGTAGTGGAAACTAGGAATT |
| Rev KRas Seq4r | GCCACTGTTTATCCAATCCAA |
| Fwd KRas amplicon 3A | CCAGTTTCTTGACTCACCTTTGAG |
| Rev KRas amplicon 3A | GGGATAAGAAAGTGCTGTGCTG |
| Fwd KRas amplicon 3B | GCCTGAAGAGAAACATAAAGAATCC |
| Rev KRas amplicon 3B | GGGAATACTGGCACTTAGAGGAA |

**human HRAS sequencing:**

| Name: | Sequence: |
|---|---|
| Fwd HRas amplicon | GCTGTGGGTTTGCCCTTCA |
| Rev HRas amplicon | CCATGTCCTGAGCTTGTGCTG |
| Fwd HRas Seq2 | GCCATCAACAACACCAAGTCTT |
| Rev HRas Seq3r | CCCATCAATGACCACCTGCTT |
| Fwd HRas Seq3 | GGCACCTGTTGGTTCTGAGTCT |
| Rev HRas Seq2r | CCCACTAAGACTCAGAACCAACA |

**human NRAS sequencing:**

| Name: | Sequence: |
|---|---|
| Fwd NRas amplicon 1 | GGGATTTCCATTGCTTAGGCT |
| Rev NRas amplicon 1 | GGTTGGGAGAGATTCAGTTGCT |
| Fwd NRas amplicon 2 | GGGACAAACCAGATAGGCAGAA |
| Rev NRas amplicon 2 | CCCAAAGCACTGACATTCCAG |
| Fwd NRas amplicon 3 | GGCTAATCTCAAACTCCTGGGTT |
| Rev NRas amplicon 3 | GGTCATTGCCAGGAATAGGGTAT |
| Fwd NRas Seq4 | CCCAGCCTGTTGTTAGGCATTT |
| Rev NRas Seq2r | GGCTGAGGCAGGAGAATCACTT |

**qPCR primers:**

| Name: | Sequence: |
|---|---|
| qRTPCR CRLF2 F | CCTTCTCCAGGAAGGTCACA |
| qRTPCR CRLLF2 R | GTCCCATTCCTGATGGAGAA |
| qRTPCR JAK2 F | CGGATAGATCACATAAAACTTCTGC |
| qRTPCR JAK2 R | TGCCAGATCCCTGTGGATA |
| mouse GAPDH qRT PCR fwd | GGTGCTGAGTATGTCGTGGA |
| mouse GAPDH qRT PCRrev | CGGAGATGATGACCCTTTTG |

**Supplementary-Tab.S3) List of oligonucleotides used to establish the mutation status of key genes involved in the analyzed pathways in the DS-ALL patients.** Forward and reverse strand oligonucleotides were used to amplify genomic regions of interests (named Amp) in PCR and all primers were then used in Sanger sequencing on produced PCR amplicons. DNA of previously(2) whole-exome-sequenced samples was used as control for both wt and mutant sequence detection.

### Supplementary Fig.S1

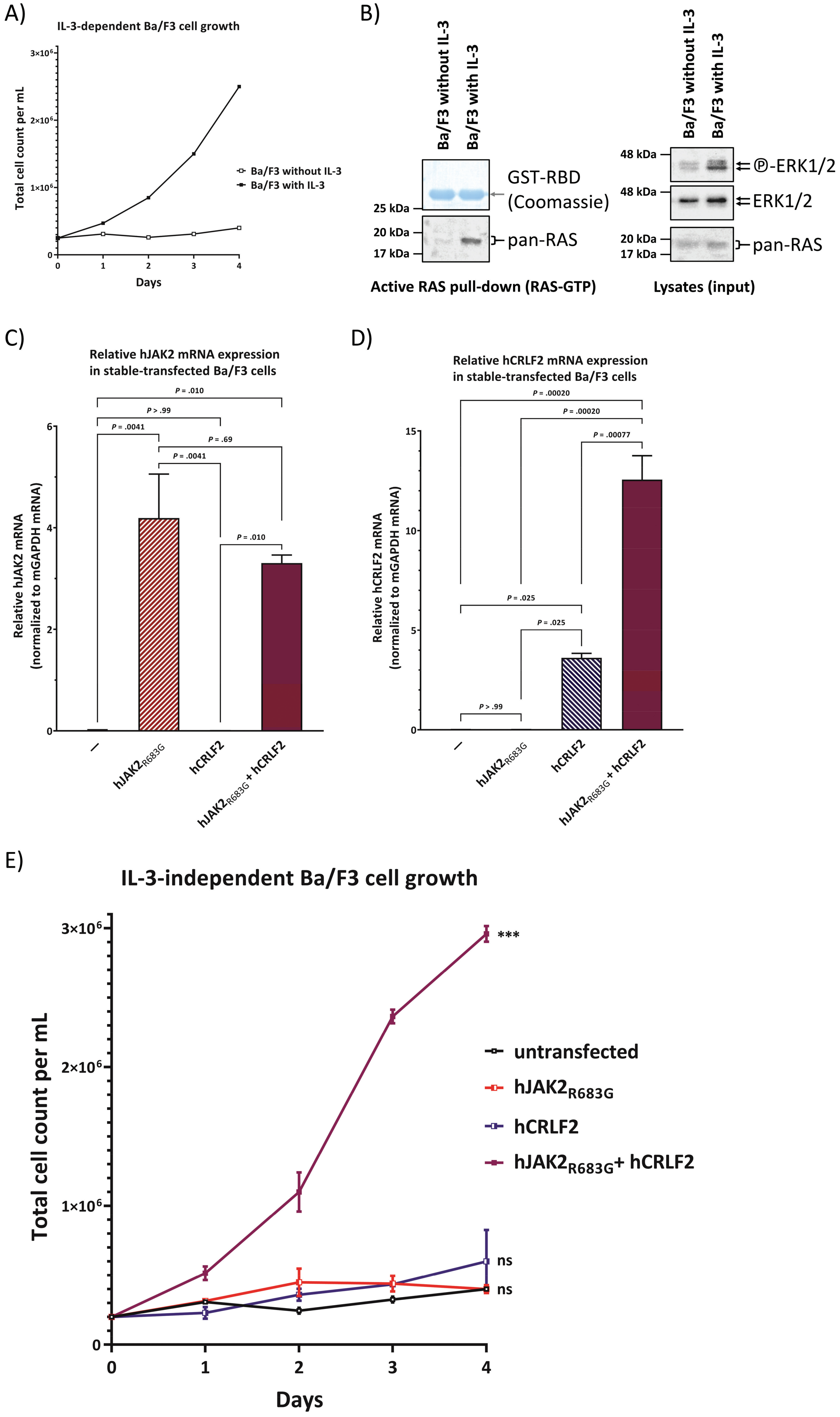

**Supplementary-Fig.S1) IL-3 induces wtRAS activity and cell growth in murine pro B cells, but can be substituted by stable overexpression of constitutively active JAK2 and CRLF2.** (A) Cell count of Ba/F3 cells (untransfected) over time. Cells were either cultured with 10 ng/mL IL-3 or without.

(B) Ba/F3 cells were cultured in 10 ng/mL IL-3. 17 hrs before lysis, the culture medium was changed to medium containing either 10 ng/mL IL-3 or no IL-3. A RAS-pull-down assay was performed. Lysates of pull-down and input were loaded on separate SDS-PAGE gels followed by Western blotting. Before Western blotting, the top part of the gel loaded with the RAS-GTP pull-down samples was stained with Coomassie dye to visualize the GST-RBD to ensure pull-down samples were loaded equally. To assess the total protein and phosphorylated protein amounts on the same PVDF-membrane, membranes were stripped and reprobed with new antibodies.

(C&D) Ba/F3 cells were stably transfected with $hJAK2_{R683G}$, hCRLF2, or both constructs together. Quantification of qPCR results for human JAK2 mRNA expression (A) and human CRLF2 mRNA expression (B) for the stably transfected Ba/F3 cell lines. All gene expression levels were normalized to murine GAPDH expression. Error bars are SD and *P*-values were determined in one-way ANOVA and post-hoc Bonferroni multiple comparison.

(E) Only the co-expression of human CRLF2 and a constitutively active $JAK2_{R683G}$ mutant construct enabled IL3-independent growth of Ba/F3 cells. The expression of either hCRLF2 or $hJAK2_{R683G}$ alone is not sufficient to promote cell growth over the course of 4 days. Data shown reproduced the experiments already published by Yoda et al.(12) to ensure that the cellular model recreated in our laboratory behaves the same and as evidence of the correlation with the state of RAS activation in these cells (Fig.1). Error bars are SD and *P*-values from a one-way ANOVA with post-hoc Bonferroni multiple comparison for the last timepoint are symbolized as asterisks (***: $P < 0.001$) or ns ($P > 0.05$).

### Supplementary Fig.S2

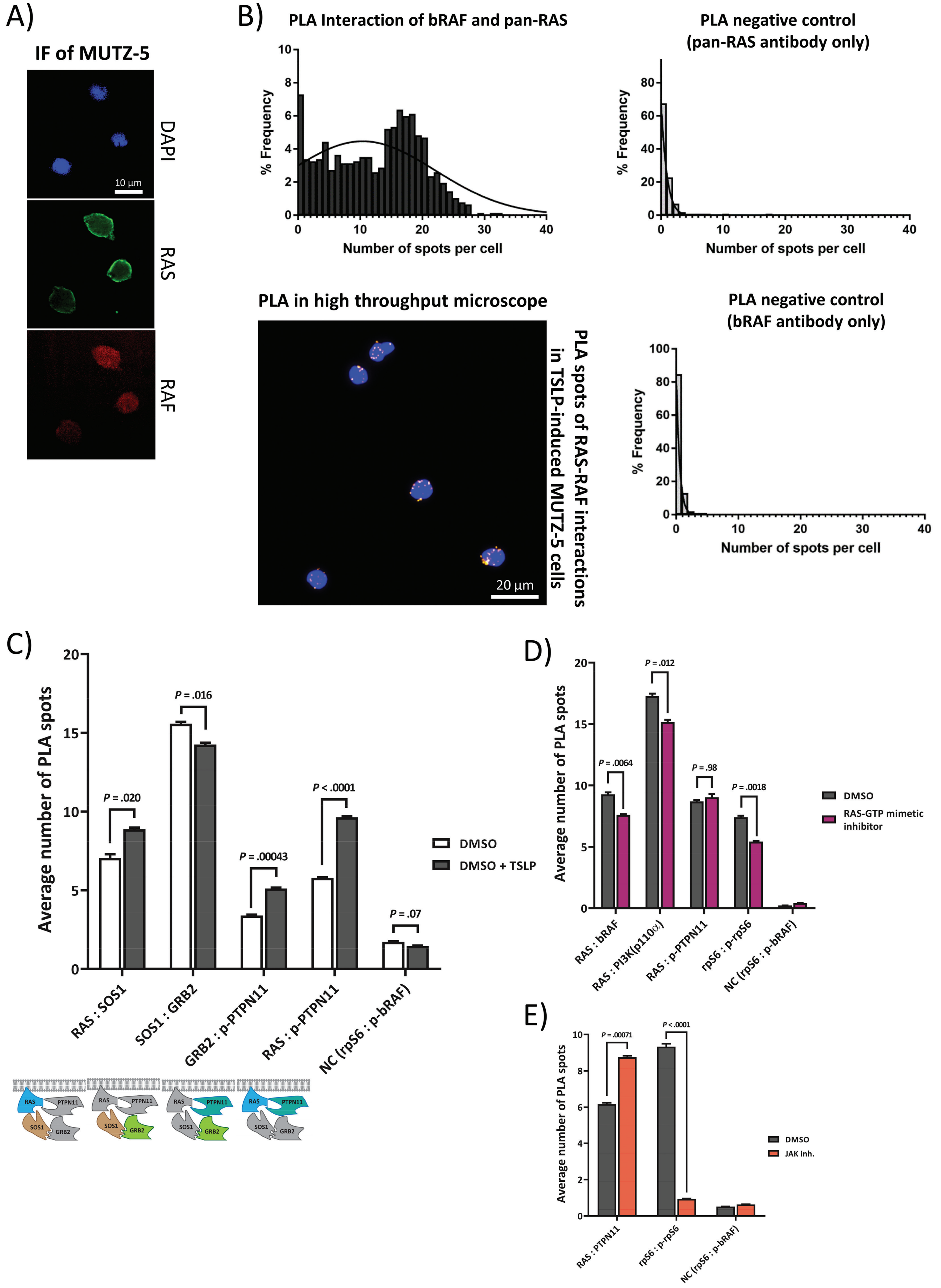

**Supplementary-Fig.S2) CRLF2-signaling promotes direct protein-protein binding (quantitated by proximity ligation assays) of multiple components involved in RAS activation in the absence of RAS mutations in human Philadelphia-like (rearranged CRLF2, JAK2$_{R683G}$) B-ALL cells.** (A) MUTZ-5 cells were PFA-fixed, permeabilized, and stained with DAPI. Primary antibodies against either pan-RAS or bRAF were used. Secondary antibodies specific to the respective primary antibody species labeled with either Alexa-488 (green) or Alexa-594 (red) were used to visualize the spatial organization of RAS (mostly plasma membrane) and RAF (mostly cytosolic) proteins in confocal microscopy.

(B) Direct interaction between individual RAS proteins and the MAPK pathway protein bRAF was monitored via proximity ligation assay (PLA) in high-throughput microscopy. MUTZ-5 cells were induced with 20 ng/mL TSLP for 5 min. After blocking, antibodies against Pan-RAS and bRAF were used in conjunction with PLA rabbit and mouse probes to allow specific readout of RAS binding to bRAF proteins in single cells in a high-throughput manner. Fluorescent microscope image shows the PLA spots in DAPI-stained cells. Histograms show the distribution of the number of spots in all cells, negative assay controls only received one of the antibodies (PLA for two non-interacting antigens showed higher levels of spots than the assay control, but remained at background levels, not shown). A minimum of 600 cells were analyzed per sample. Non-linear Gaussian fitting curves were plotted.

(C) Direct interaction between protein pairs involved in RAS activation (see cartoon below the graph) was monitored via PLA in high-throughput microscopy. MUTZ-5 cells were either not induced or induced with 20 ng/mL TSLP for 10 min. Cells were fixed and permeabilized in a 96 well plate. After blocking, the indicated antibody pairs were used in conjunction with PLA rabbit and mouse probes to allow the amplification and staining of interaction-specific PLA spots. The negative control (NC) samples used antibodies for two cytosolic proteins (rpS6 and phosphorylated bRAF) that were not expected to directly interact. At least 1900 cells per condition were analyzed in a high-throughput manner. The bar graph shows the averages of 3 technical replicates. Error bars are SD and *P*-values shown are Student's T-test after Bonferroni correction for sequential multiple comparison for all uninduced vs TSLP-induced pairs.

(D) Direct interaction between protein pairs involved in RAS activation and the binding between RAS to RAS effectors was monitored via PLA in high-throughput microscopy. MUTZ-5 cells treated with 20 ng/mL TSLP were pre-treated with either DMSO or the RAS-GTP mimetic inhibitor Rigosertib (30 µM) for 1.5 hrs. Cells were fixed and permeabilized in a 96 well plate. After blocking, the indicated antibody pairs were used in conjunction with PLA rabbit and mouse probes to allow the amplification and staining of interaction-specific PLA spots. The negative control (NC) samples used antibodies for two cytosolic proteins (rpS6 and phosphorylated bRAF) that were not expected to directly interact. At least 700 cells per condition were analyzed in a high-throughput manner. The bar graph shows the averages of 3 technical replicates. Error bars are SD and *P*-values shown are Student's T-test after Bonferroni correction for sequential multiple comparison for all DMSO vs Rigosertib pairs.

(E) Direct interaction between protein pairs between RAS and PTPN11 was measured via PLA in high-throughput microscopy. MUTZ-5 cells treated with 20 ng/mL TSLP were pre-treated with either DMSO or 5 µM Ruxolitinib (JAK inh.) for 1.5 hrs. Cells were handled like in (D). The condition using antibodies for rpS6 and phosphorylated rpS6 verified that the JAK inhibitor was effective in this experiment. At least 20,000 cells per condition were analyzed in a high-throughput manner. The bar graph shows the averages of 3 technical replicates. Error bars are SD and *P*-values shown are Student's T-test after Bonferroni correction for sequential multiple comparison for all DMSO vs Ruxolitinib pairs.

Supplementary Fig.S3

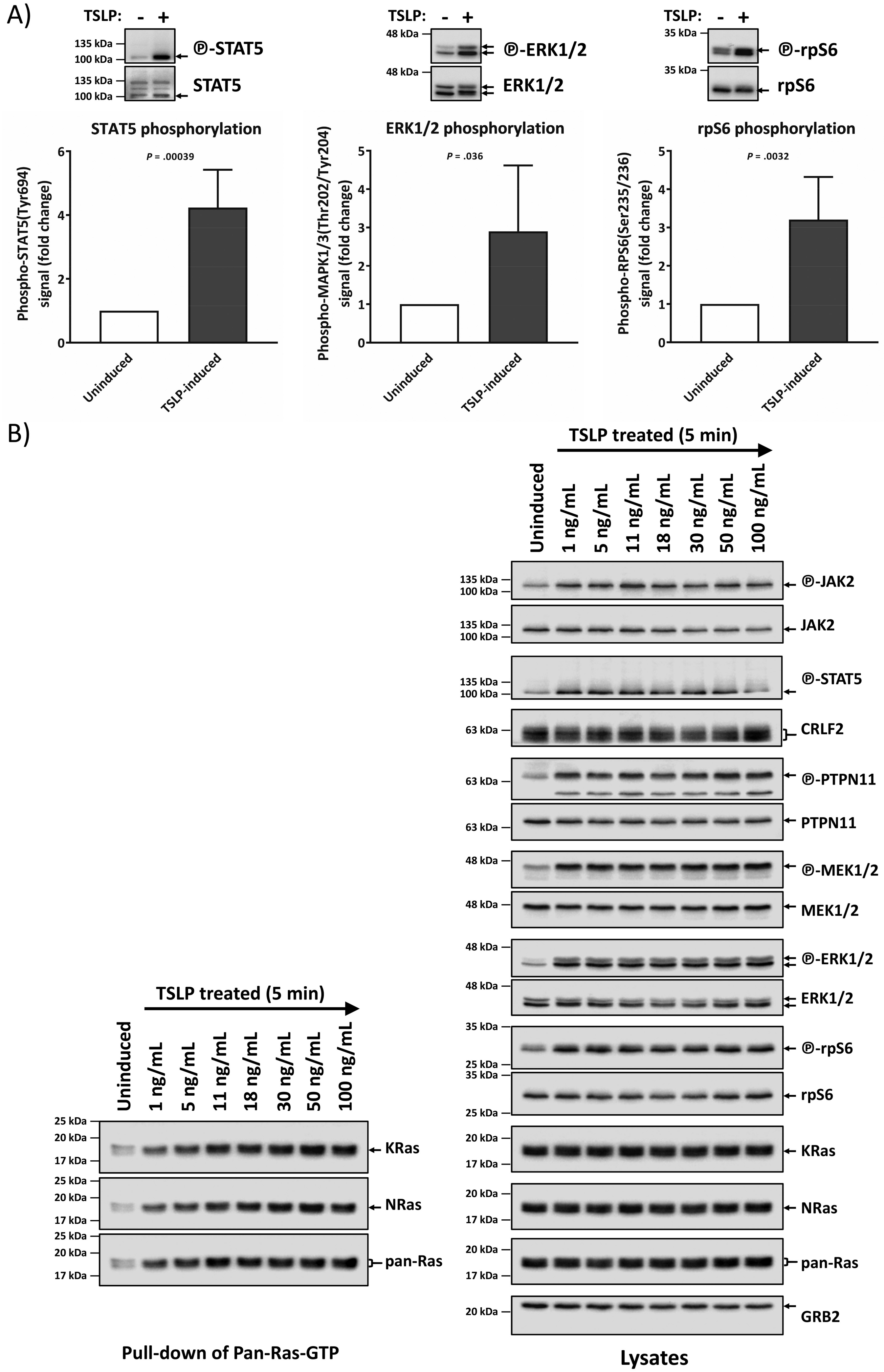

**Supplementary-Fig.S3) TSLP-induction actives STAT and PI3K/mTOR signaling in Ph-like B ALL cells. Titration of TSLP concentration in Ph-like B ALL cells.** (A) Extension of Fig.2A. MUTZ-5 cells (Human Ph-like B-ALL cells bearing CRLF2-rearranged and spontaneous JAK2R683G mutation) were stimulated (or not) with 20 ng/mL human TSLP for 10 min before cell lysis. Lysates were loaded on an SDS-PAGE gel followed by Western blotting. To assess the total protein and phosphorylated protein amounts on the same PVDF-membrane, membranes were stripped and reprobed with new antibodies. The experiment was repeated 5 times independently and the graphs show the quantification for active STAT5 (phosphorylated STAT5), active ERK1/2 (phosphorylated ERK1/2) and active RPS6 (phosphorylated RPS6). Representative Western blots above the respective graph show the phosphorylated protein form and the total expression of each protein.

(B) Effect of TSLP concentration titration. MUTZ-5 cells were incubated with the indicated amounts of TSLP (0 ng/mL to 100 ng/mL) for 5 min and then the cells were lysed on ice. Each cell lysate was split up for analysis in RAS-GTP pull-down assay and for total protein signal. RAS-GTP pull-down and lysate samples were loaded on separate gels. An SDS-PAGE followed by Western blotting was performed. To assess the total protein and phosphorylated protein amounts on the same PVDF-membrane, membranes were stripped and reprobed with new antibodies. Left-hand side blots show the RAS-GTP pull-down while the right-hand side blots show whole cell lysates of the same samples. Antibody-targets are labeled on the right side of each image with black arrows indicating the respective protein band.

Supplementary Fig.S4

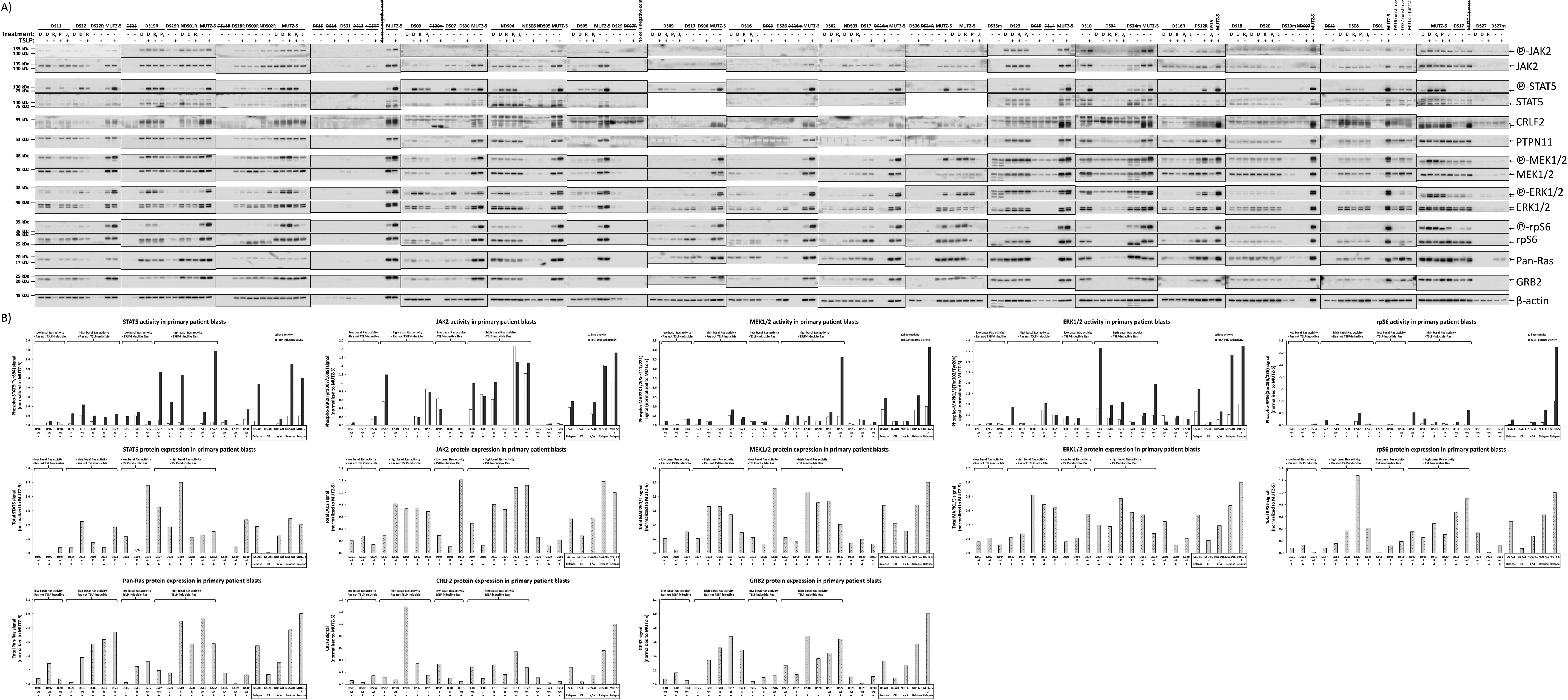

**Supplementary-Fig.S4) Western blots for all analyzed patient samples and the resulting quantified data for the sub-stratification of DS-ALL by PCA.** (A) Primary samples of DS-ALL patients at presentation, relapse, or remission, as well as MUTZ-5 cells, were thawed and cells were gently recovered in 9 mL RPMI1640 (Cat.#11-875-119; Life Technologies, Carlsbad, US) containing 20% characterized fetal bovine serum (FBS; Cat.#SH30071.03; GE Healthcare, Chicago, US), 20 U/mL Benzonase (Cat.#70746; EMD Millipore), 2 mM L-Glutamine (Cat.#25030081 Life Technologies), and 100 U/mL Penicillin-Streptomycin (Cat.#15140122, Life Technologies). Cells were resuspended in 5 mL fresh medium without Benzonase. Cells were counted using the automated cell-counter NucleoCounter NC-250 (ChemoMetec, Allerod, DK) and viability was assessed via staining with Solution18 (Cat.#910-3018, ChemoMetec). Cells were seeded at $1.5\times10^6$ cells/mL density in IMDM-complete (IMDM (Cat.#12440053; Life Technologies) with 10% FBS, 2 mM L-Glutamine, 100 U/mL Penicillin-Streptomycin, 200 µg/mL apo-Transferrin (Cat.#11096-37-0; Santa Cruz Biotechnlogy), 0.1% 2-Mercaptoethanol (Cat.#21985023; Life Technologies), 1 µg/mL insulin (Cat.#11061-68-0; Sigma-Aldrich), 10 ng/mL IL-3 (Cat.#200-03; Peprotech, Rocky Hill, US) and 10 ng/mL IL-7 (Cat.#200-07; Peprotech). After 24 hrs, surviving cells were reseeded at $1\times10^6$ viable cells/mL density in IMDM-complete. After 16 hrs the cells were reseeded at $1\times10^6$ cells/mL density in 2 mL OptiMEM (Cat.#31985070; Life Technologies) with 5% FBS. After 3 hrs, cells were either left uninduced or induced with 20 ng/mL TSLP (Cat.#1398-TS; R&D Systems) for 10 min at 37 °C before lysis. Where indicated, patient cells were pre-treated during the 3 hrs incubation with DMSO (D), RAS-inhibitor ($R_i$), PI3K/mTOR inhibitor ($P_i$), or JAK inhibitor ($J_i$). For Western blot analysis, cells were lysed and loaded on an SDS-PAGE gel. To assess the total protein and phosphorylated protein amounts on the same PVDF-membrane, membranes were stripped and reprobed with new antibodies. Antibody-targets are labeled on the right side of each image with black arrows indicating the respective protein band. As all lanes are shown for all gels in every specific staining, sample names that are crossed-out were either not part of the analysis, or showed too low loading in all expressed proteins to be used for any quantification. For blot imaging, each PVDF membrane piece was separately incubated with chemiluminescent HRP substrate solution (Cat.#WBKLS0500; EMD Millipore) for 2 min and imaged via the auto-exposure function of the ChemiDoc MP imaging system (Bio-Rad Laboratories) within 100 sec exposure time to standardize the sensitive but otherwise semi-quantifiable enzymatic-based detection.

(B) Quantification of Western blot signals in (A) for all DS-ALL samples at presentation as well as Non-DS (NDS) at presentation, DS complete remission (CR), and DS/NDS at relapse (boxed group at right end of each bar graph). For quantification, the raw images were analyzed in Fiji 1.52n (ImageJ, National Institutes of Health, US) using the subtract background process followed by measuring the signal peaks in the Gel Analyzer function. Microsoft Excel software was used to adjust all samples to either their loading control (β-actin) or the quantitative total protein fluorescent signal (AzureRed, Cat.#AC2124; Azure Biosystems, Dublin, US). All samples' protein activity and total protein signals were normalized to the respective signal from the uninduced MUTZ-5 samples loaded as reference on each membrane. 81% of all analyzed samples showed sufficient signal in loading control and tested proteins to be appropriate for further normalization and analysis. The quantified phosphorylated protein signals were additionally adjusted to the respective quantified total protein loading. Brackets on top indicate the groups of the four RAS activity patterns presented in Fig.3C. White and black bar graphs show the basal and TSLP-induced activation levels, respectively, of STAT5, JAK2, MEK1/2, ERK1/2, and RPS6 while the dotted grey bar graphs show the respective total protein expression levels. Outcome of leukemia is symbolized as circle for good outcome and triangle for poor outcome; RAS pathway mutations (R), JAK2 mutations (J), or neither (wt) are listed for each DS-ALL patient.

### Supplementary Fig.S5

A)

| Field | PC1 | PC2 | PC3 |
|---|---|---|---|
| Dispers | 0.385601 | 0.049523 | 0.081659 |
| Overall | 0.385601 | 0.435125 | 0.516784 |
| Pan-Ras basal activity | 0.214594 | **0.485179** | 0.048106 |
| Pan-Ras activity with TSLP | 0.227889 | **0.474698** | 0.041787 |
| JAK2 basal activity | 0.237394 | -0.24357 | -0.40745 |
| JAK2 activity with TSLP | 0.297418 | -0.1813 | -0.35807 |
| STAT5 basal activity | 0.172951 | -0.20932 | 0.065401 |
| STAT5 activity with TSLP | **0.322827** | 0.163583 | -0.23887 |
| MEK basal activity | 0.311386 | 0.309735 | 0.091045 |
| MEK activity with TSLP | **0.351922** | 0.1494 | 0.076544 |
| ERK basal activity | 0.281839 | -0.06334 | -0.04313 |
| ERK activity with TSLP | 0.287373 | -0.22426 | -0.26412 |
| S6 basal activity | 0.226173 | -0.24303 | **0.609553** |
| S6 activity with TSLP | 0.280702 | -0.32844 | **0.303667** |
| CRLF2 protein expression | 0.250139 | -0.18761 | 0.007298 |

D)

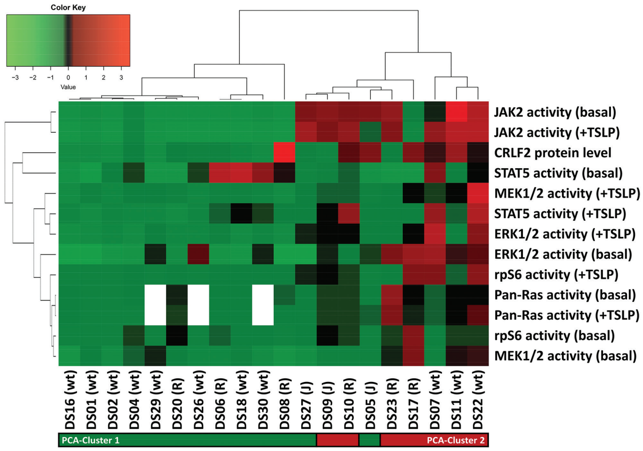

B)

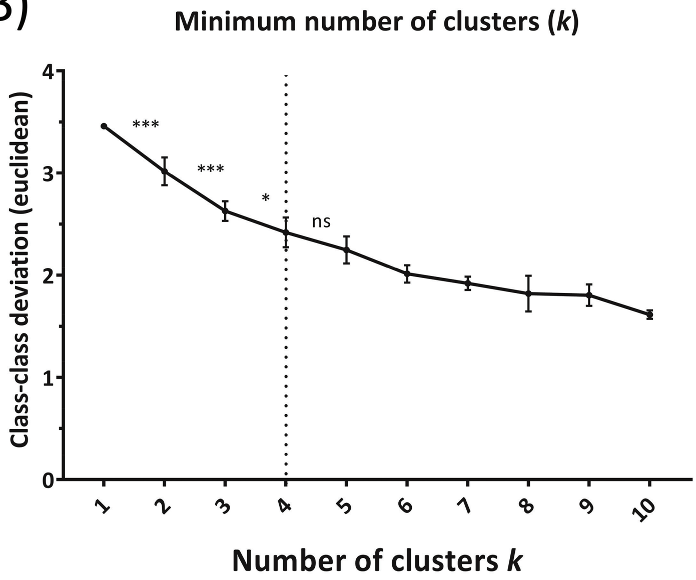

C)

| | N = | DS patients with good outcome | DS patients with poor outcome | DS relapse samples | DS Complete remission sample | DS Remission sample with imminent relapse | NDS patients with good outcome | NDS patients with poor outcome | NDS relapse samples | NDS (Ph-like) relapse cell line (MUTZ-5) |
|---|---|---|---|---|---|---|---|---|---|---|
| **Total** | **38** | **11** | **9** | **7** | **2** | **2** | **3** | **1** | **2** | **1** |
| Cluster 1 | 19 | 10 | 3 | 1 | 2 | 0 | 2 | 1 | 0 | 0 |
| Cluster 2 | 15 | 1 | 6 | 4 | 0 | 2 | 1 | 0 | 1 | 0 |
| Cluster 3 | 3 | 0 | 0 | 1 | 0 | 0 | 0 | 0 | 1 | 1 |
| Cluster 4 | 1 | 0 | 0 | 1 | 0 | 0 | 0 | 0 | 0 | 0 |

**Supplementary-Fig.S5) PCA statistics and PCA cluster counts on primary DS-ALL and NDS-ALL blast cells.** (A) Basal and induced activation levels for RAS, JAK2, STAT5, MEK, ERK, and RPS6, as well as CRLF2 protein expression, were fed into ViDaExpert v1.2 to calculate 3 main principle components (PC1-3) in a PCA analysis. Numbers in bold highlight the categories (field) with the greatest correlation within each PC.

(B) Euclidean *k*-means clustering was performed in ViDaExpert v1.2 on the PCA analysis in (A) for $k$ = 1 to 10 (6 times each). In order to identify the minimal number of clusters needed for the grouping analysis, class-class deviation was averaged for each k (error bars are SD). Class-class deviation from sequential, increasing number of clusters stopped being statistically significant at $k$ = 4 (doted line); * ($P < .05$), *** ($P < .001$), ns (not significant).

(C) Number of samples within each of the four clusters identified in the *k*-means clustering of the PCA data represented in Fig.4A. Count of the different patient sample categories for each cluster.

(D) Unsupervised hierarchical clustering of all analyzed DS-ALL presentation samples for all 6 measured protein activities (basal and TSLP-induced) as well as CRLF2 protein expression. The clustering results were similar to the independent *K*-means clustering (indicated by the red/green colored bar underneath the sample IDs) of the PCA (Fig.4A), with 90% (18/20) of the samples clustered in the same groups by both methods. Presence of RAS pathway mutations (R), JAK2 mutations (J), or neither (wt) is indicated (no single patient in this cohort was found to have both RAS and JAK2 mutations).

Supplementary Fig.S6

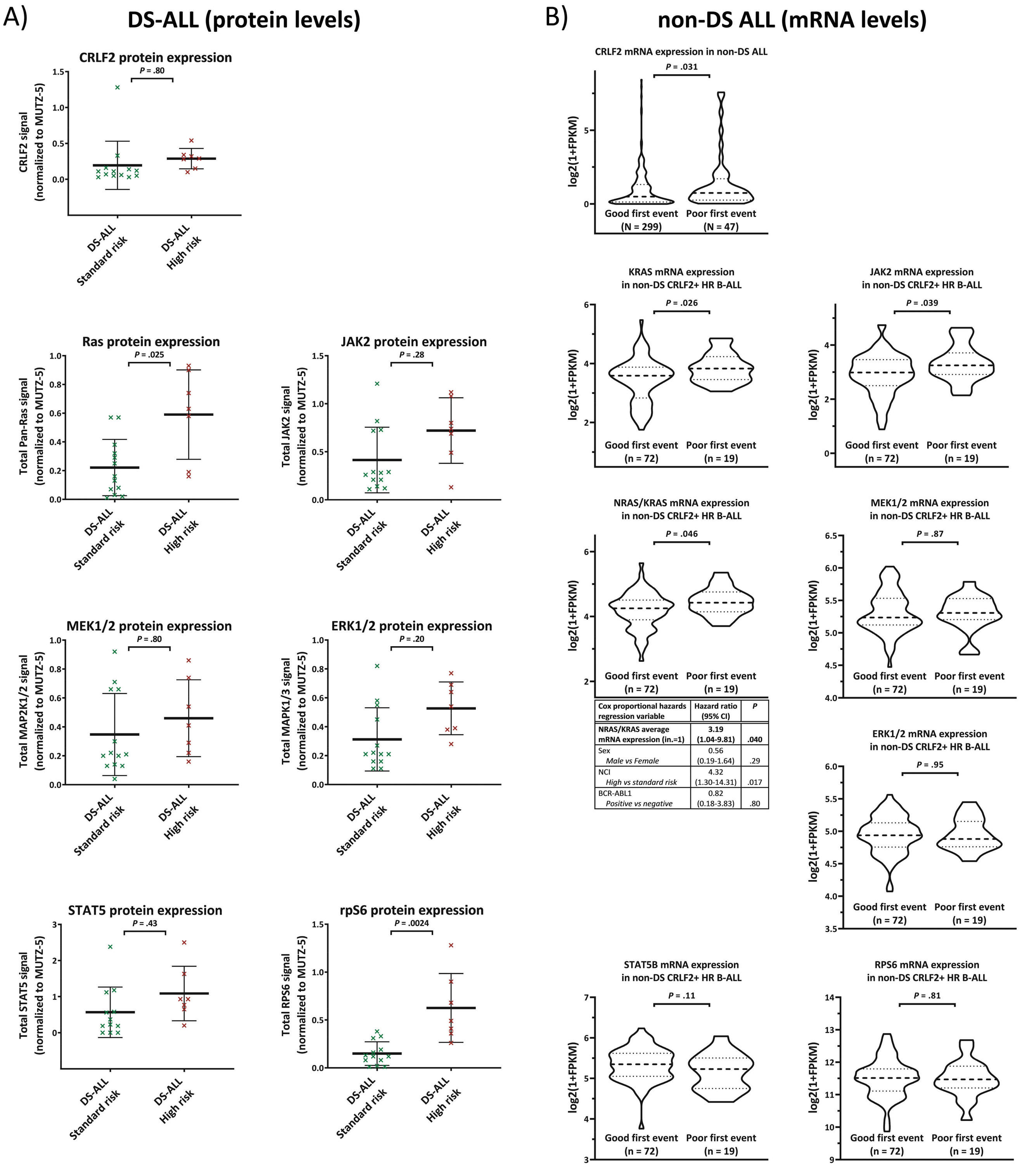

| Cox proportional hazards regression variable | Hazard ratio (95% CI) | P |
|---|---|---|
| **NRAS/KRAS average mRNA expression (in.=1)** | **3.19 (1.04-9.81)** | **.040** |
| Sex *Male vs Female* | 0.56 (0.19-1.64) | .29 |
| NCI *High vs standard risk* | 4.32 (1.30-14.31) | .017 |
| BCR-ABL1 *Positive vs negative* | 0.82 (0.18-3.83) | .80 |

**Supplementary-Fig.S6) Disease outcome-correlations with expression levels of components used in RAS-pathway sub-stratification. Protein levels in DS-ALL (A) and mRNA levels in non-DS childhood ALL (MS2003/2010 cohorts) (B)**

(A) Average protein expressions of analyzed key pathway components are compared between the SR group (DS-ALL patients at diagnosis in PCA cluster 1) and the HR group (DS-ALL patients at diagnosis in PCA cluster 2) identified in Fig.4. All error bars are SD. *P*-values shown are Student's T-test after Bonferroni correction for sequential multiple comparison for all proteins analyzed.

(B) Violin blots of the mRNA levels based on whole transcriptome RNAseq of the MS2003/2010 cohorts at diagnosis are shown for gene(s) equivalents to proteins analyzed in the DS-ALL cohort. The dashed line represents the median while the dotted lines above and below the median mark the respective quartiles. *CRLF2* was analyzed for outcome on the total non-DS ALL cohort (N = 346). Subsequently, all other shown mRNA expression levels (*KRAS*, *KRAS*&*NRAS* combined, *STAT5B*, *JAK2*, *MEK1*&*2* combined, *ERK1*&*2* combined, *RPS6*) were compared in B-ALL samples positive for *CRLF2*-mRNA expression (log2(1+FPKM) > 0.7) for the first event outcome. Poor first event for the MS2003/2010 cohort was defined as resistance, death or relapse; good first event means complete remission. Subtypes that are known to favor a good outcome were excluded from the analysis (ETV6-RUNX1, hyperdiploid, DUX4 and ZNF384). The resulting n=91 subcohort harbored 13 *RAS* mutations (4/19 poor outcome samples) and 6 *JAK*2 mutations (0/19 poor outcome patients) but never both together. For *KRAS*&*NRAS* combined, table details results of Cox proportional hazards regression. All error bars are SD. *P*-values shown were calculated using Student's T-test. Bonferroni-*P*-values adjusted for sequential multiple comparison were not ≤ α (0.05) and are listed in Supplementary-Tab.S2.

### Supplementary Fig.S7

#### A) Non-DS ALL mRNA levels (full MS2003/2010 cohort)

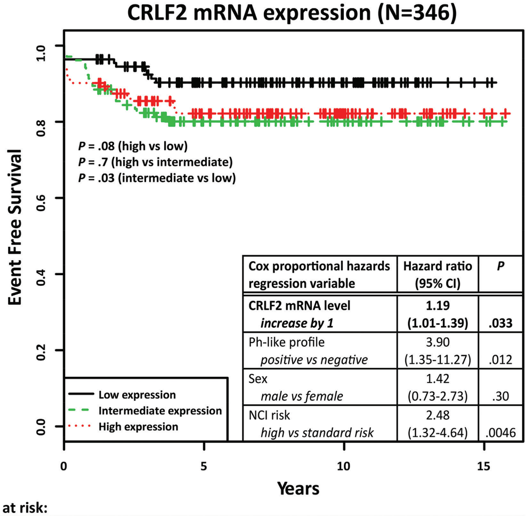

| Cox proportional hazards regression variable | Hazard ratio (95% CI) | *P* |
|---|---|---|
| **CRLF2 mRNA level** ***increase by 1*** | **1.19 (1.01-1.39)** | **.033** |
| Ph-like profile *positive vs negative* | 3.90 (1.35-11.27) | .012 |
| Sex *male vs female* | 1.42 (0.73-2.73) | .30 |
| NCI risk *high vs standard risk* | 2.48 (1.32-4.64) | .0046 |

| Number at risk: | | | | | | | | | | | | | | | | |
|---|---|---|---|---|---|---|---|---|---|---|---|---|---|---|---|---|
| Low expression | 111 | 107 | 98 | 89 | 83 | 71 | 65 | 58 | 47 | 40 | 33 | 24 | 14 | 6 | 4 | 2 |
| Interm. expression | 104 | 94 | 84 | 78 | 63 | 55 | 51 | 44 | 37 | 30 | 26 | 16 | 15 | 10 | 5 | 3 |
| High expression | 112 | 101 | 92 | 84 | 76 | 70 | 66 | 57 | 47 | 44 | 31 | 23 | 18 | 12 | 4 | 2 |

#### B) Non-DS ALL mRNA levels (CRLF2+, high-risk genetics subcohort)

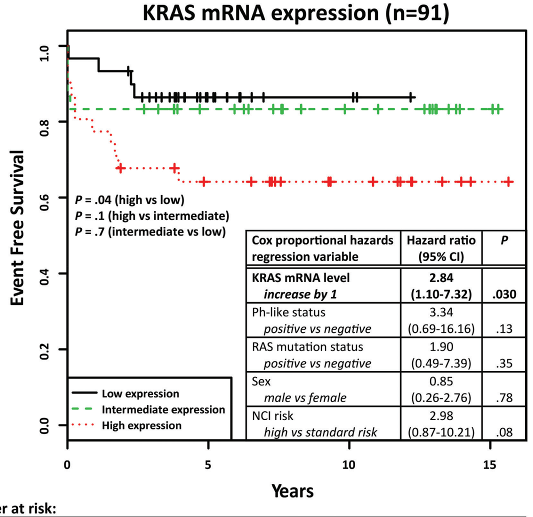

| Cox proportional hazards regression variable | Hazard ratio (95% CI) | *P* |
|---|---|---|
| **KRAS mRNA level** ***increase by 1*** | **2.84 (1.10-7.32)** | **.030** |
| Ph-like status *positive vs negative* | 3.34 (0.69-16.16) | .13 |
| RAS mutation status *positive vs negative* | 1.90 (0.49-7.39) | .35 |
| Sex *male vs female* | 0.85 (0.26-2.76) | .78 |
| NCI risk *high vs standard risk* | 2.98 (0.87-10.21) | .08 |

| Number at risk: | | | | | | | | | | | | | | | | |
|---|---|---|---|---|---|---|---|---|---|---|---|---|---|---|---|---|
| Low expression | 30 | 29 | 28 | 23 | 17 | 11 | 7 | 3 | 3 | 3 | 3 | 1 | 1 | 0 | 0 | 0 |
| Interm. expression | 30 | 25 | 25 | 24 | 20 | 19 | 18 | 16 | 13 | 12 | 11 | 11 | 10 | 7 | 2 | 2 |
| High expression | 31 | 24 | 20 | 20 | 18 | 17 | 17 | 16 | 12 | 12 | 9 | 8 | 6 | 4 | 2 | 1 |

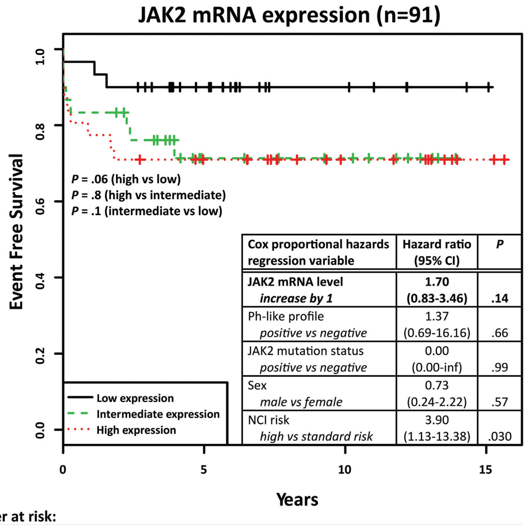

| Cox proportional hazards regression variable | Hazard ratio (95% CI) | *P* |
|---|---|---|
| **JAK2 mRNA level** ***increase by 1*** | **1.70 (0.83-3.46)** | **.14** |
| Ph-like profile *positive vs negative* | 1.37 (0.69-16.16) | .66 |
| JAK2 mutation status *positive vs negative* | 0.00 (0.00-inf) | .99 |
| Sex *male vs female* | 0.73 (0.24-2.22) | .57 |
| NCI risk *high vs standard risk* | 3.90 (1.13-13.38) | .030 |

| Number at risk: | | | | | | | | | | | | | | | | |
|---|---|---|---|---|---|---|---|---|---|---|---|---|---|---|---|---|
| Low expression | 30 | 29 | 27 | 25 | 19 | 17 | 12 | 8 | 6 | 6 | 6 | 5 | 4 | 2 | 2 | 1 |
| Interm. expression | 30 | 25 | 24 | 21 | 15 | 11 | 11 | 10 | 9 | 9 | 8 | 6 | 5 | 3 | 0 | 0 |
| High expression | 31 | 24 | 22 | 21 | 21 | 19 | 19 | 17 | 13 | 12 | 9 | 9 | 8 | 6 | 2 | 2 |

**Supplementary-Fig.S7) Disease outcome-correlations and multivariate analysis for mRNA expression levels in non-DS childhood ALL (MS2003/2010 cohorts) of key components emanating from RAS-pathway sub-stratification in primary samples at diagnosis** (A) Kaplan–Meier estimates of event-free survival of all non-DS ALL patients (N=346) according to *CRLF2*-mRNA expression. Low, intermediate, and high curves for *CRLF2*-mRNA expression levels (overall P = .09) each represent one third of the cohort (bottom, mid, and top third, respectively). Table within the graph shows multivariate analysis using Cox proportional-hazards model for *CRLF2*-mRNA levels together with the prognostic factors Ph-like status, sex, and NCI risk groups. Patient numbers at risk for each year are given in the table below the survival curve.

(B) Subsequently, all other mRNA expression levels (*KRAS*, JAK2, *KRAS*&*NRAS* combined, *STAT5B*, *MEK1*&*2* combined, *ERK1*&*2* combined, *RPS6*) were compared in B-ALL samples positive for *CRLF2*-mRNA expression (log2(1+FPKM) > 0.7) for the first event outcome (Supplementary-Fig.S6B). Subtypes that are known to favor a good outcome were excluded from the analysis (ETV6-RUNX1, hyperdiploid, TCF3-PBX1, DUX4, and ZNF384). The resulting n=91 subcohort harbored 13 *RAS* mutations (4/19 poor outcome samples) and 6 *JAK2* mutations (0/19 poor outcome patients), but never both together. Kaplan–Meier curves show estimates of event-free survival of the HR non-DS ALL subcohort (n=91) grouped by *RAS* (overall P = .08) or *JAK2*-mRNA (overall P = .2) expression levels. The tables within the graphs show multivariate analysis using Cox proportional-hazards model for either *RAS*-mRNA or *JAK2*-mRNA levels respectively, together with *KRAS/NRAS* or *JAK2* activating mutations respectively, and the prognostic factors Ph-like status, sex, and NCI risk groups.

### Supplementary Fig.S8

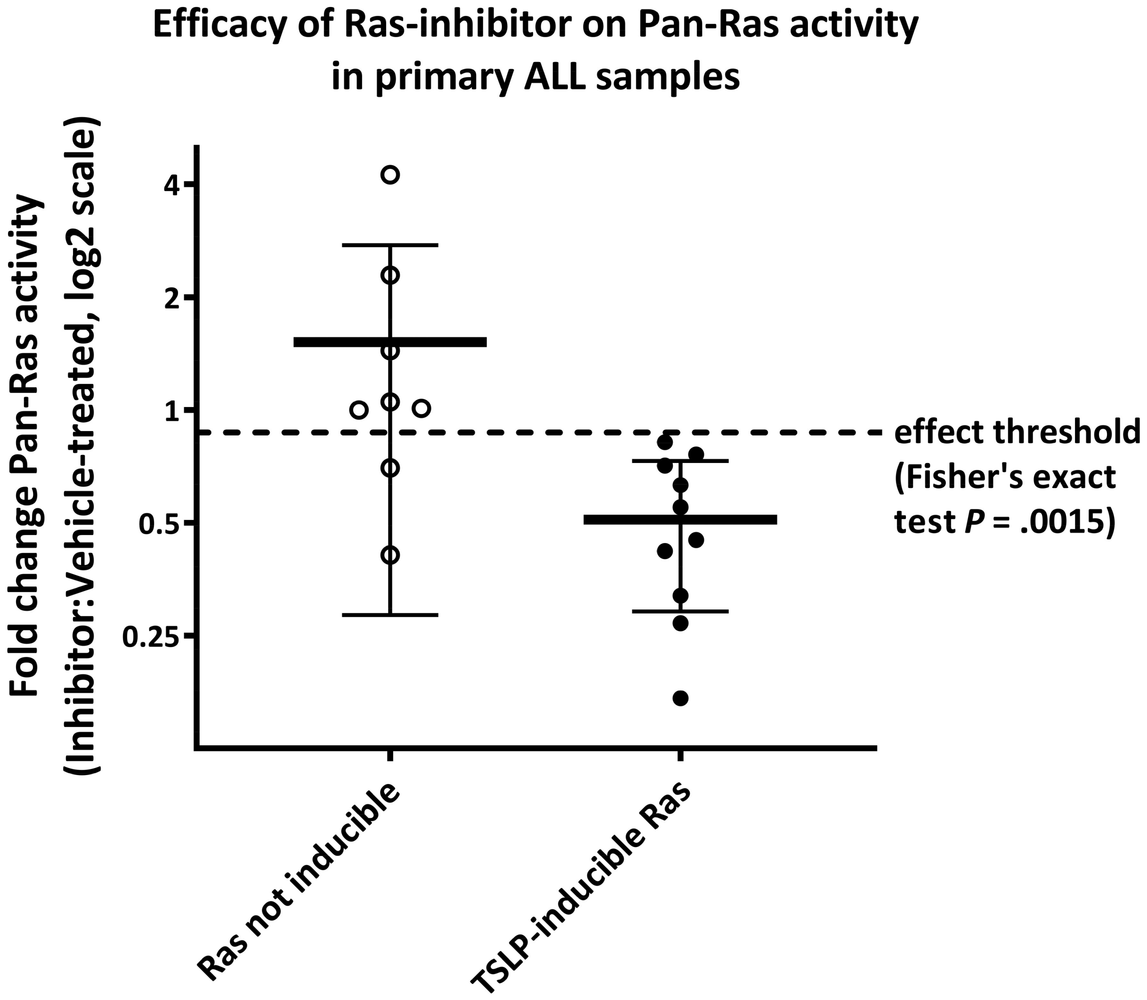

**Supplementary-Fig.S8) Efficacy of the RAS inhibitor stratified by patient sample inductility.** Primary presentation samples of DS-ALL and non-DS ALL patients (characterized in Supplementary-Tab.S1) were cultured for 2 days (see Supplementary-Fig.S4A for details). Samples with sufficient cell count were treated with either 0.5% DMSO (vehicle control) or 50 µM Salirasib (indirect pan-RAS inh.) for 3 hrs after which the cells were induced for 10 min with 20 ng/mL TSLP in serum-reduced medium. Cells were lysed and a RAS-GTP ELISA pull-down assay was performed. The efficacy of the RAS inhibitor (expressed as RAS activity of inhibitor-treated divided by RAS activity of DMSO-treated) on ELISA-measured RAS activity in patient samples that were defined in Fig.3B as not TSLP-inducible for RAS is compared to those in which RAS activity was inducible by TSLP. If inhibitor treatment reduced the RAS activity by over 10% compared to vehicle-control (dashed line in plot), the sample was tallied as successful RAS blocking. A Fisher's exact test was performed between the groups.
